# NuSAP participates in metaphase spindle length control in mammalians

**DOI:** 10.1101/2023.02.26.530101

**Authors:** Yao Wang, Mengjie Sun, Guangwei Xin, Biying Yang, Qing Jiang, Chuanmao Zhang

## Abstract

Precise chromosome congression and segregation require proper assembly of a steady-state metaphase spindle, which is dynamic and maintained by continuous microtubule flux. NuSAP is a microtubule-stabilizing and -bundling protein that promotes chromosome-dependent spindle assembly. However, its function in spindle dynamics remains unclear. Here, we demonstrate that NuSAP regulates the dynamics and length control of the metaphase spindle. Mechanistically, NuSAP facilitates kinetochore capture and spindle assembly via promoting Eg5 binding with microtubules. It also prevents excessive microtubule depolymerization through interacting with Kif2A and reduces its spindle-pole localization. NuSAP is phosphorylated by Aurora A at Ser-240 during mitosis, and this phosphorylation promotes its interaction with Kif2A on the spindle body and reduces its localization to the spindle poles, thus maintaining the proper spindle microtubule flux. NuSAP knockout resulted in shorter spindle formation with faster microtubule flux and chromosome misalignment. Taken together, we uncover that NuSAP participates in spindle assembly, dynamics, and metaphase spindle length control via affecting microtubule flux and Kif2A localization.

## Introduction

The mitotic spindle is a well-organized and evolutionarily conserved microtubule (MT)-based structure that mediates precise chromosome congression and segregation (Karsenti & Vernos, 2001; Mitchison & Salmon, 2001). The spindle MTs grow and shrink by addition and loss of tubulin subunits at their ends, allowing the spindle to efficiently probe the cytoplasmic space for biorientation of sister kinetochores and to congress chromosomes to the metaphase plate (Gadde & Heald, 2004; Kline-Smith & Walczak, 2004). Although the spindle MTs are dynamically unstable and switch between growing and shrinking states, the metaphase spindle will finally reach to a balance that maintains a constant length and well-constructed structure (Dumont & Mitchison, 2009; Rogers, Rogers, & Sharp, 2005). In addition, mitotic spindle MTs display poleward flux, which is driven by four kinesins. Accordingly, interpolar MTs are slid apart by MT-sliding motors Eg5 and Kif15, sequentially contributed by CENP-E at KTs in prometaphase and Kif4A on chromosome arms in metaphase (Steblyanko et al., 2020). When outward sliding apart of interpolar MTs is balanced by polymerization and depolymerization of the spindle MTs at the minus-ends and plus-ends, the spindle maintains a steady length (Brust-Mascher, Sommi, Cheerambathur, & Scholey, 2009; Fink et al., 2009; Goshima & Scholey, 2010; Steblyanko et al., 2020). This balance is achieved mainly via kinesin-13 family proteins (Brust-Mascher & Scholey, 2002; Brust-Mascher et al., 2009; Ems-McClung & Walczak, 2010; Ganem & Compton, 2004; Miyamoto, Perlman, Burbank, Groen, & Mitchison, 2004; Saunders, Powers, Strome, & Saxton, 2007), in concert with other regulators of the spindle MT dynamics (Buster, Zhang, & Sharp, 2007; Goshima & Scholey, 2010; Kwok & Kapoor, 2007).

NuSAP (nucleolar and spindle-associated protein) is a conservative and approximately 55 kDa protein, and is selectively expressed in proliferating cells (Raemaekers et al., 2003). It contains a putative DNA-binding motif in its N-terminal part, a nuclear localization sequence in the N-terminal part near to the middle of the molecule, a KEN-box in its C-terminal part that serves as a general targeting signal for the anaphase-promoting complex/cyclosome (APC/C)-Cdh1 E3 ubiquitin ligase, a potential PEST sequence at its C-terminal part for protein degradation, and a microtubule-binding domain at the C-terminal part (Iyer, Moghe, Furukawa, & Tsai, 2011; L. Li et al., 2007; Raemaekers et al., 2003; Ribbeck et al., 2006). In interphase, NuSAP is nucleolar but shuttles between the nucleus and the cytoplasm in an importins-dependent manner(Ribbeck et al., 2006). In mitosis, NuSAP is firstly localized to the chromosomes and then to the chromosome-proximal MTs, to the central spindle MTs and finally to the midbody along with the cell division cycle progression. NuSAP is an essential protein for individual development. Its knockout in mouse is early embryonic lethality (Vanden Bosch et al., 2010), and in zebrafish impairs migration of neural crest cells (Nie, Wang, He, & Huang, 2010). Abnormal expression of NuSAP also links to many sorts of cancers (Iyer et al., 2011).

The precise functions of NuSAP in cells remain largely unclear, especially in interphase; but, as a microtubule-associated protein (MAP), it plays an important role in chromosome-dependent spindle assembly (Raemaekers et al., 2003; Ribbeck et al., 2006; Ribbeck, Raemaekers, Carmeliet, & Mattaj, 2007). Previous studies suggested that NuSAP is a mitotic Ran GTPase target that stabilizes and cross-links MTs (Ribbeck et al., 2006). During mitosis, NuSAP is dissociated with importins and immobilized on chromatin to induce MTs attachment to the chromosomes (Ribbeck et al., 2007). Both overexpression and knockdown of NuSAP resulted in defective mitotic spindle formation and chromosome segregation (Raemaekers et al., 2003). Despite the efforts to explore NuSAP roles in mitotic spindle assembly, its function in spindle dynamics remains unclear.

In this work, we reveal that NuSAP regulates the spindle dynamics to maintain metaphase spindle length through coordinating MT depolymerization and MT-MT sliding. We find that NuSAP is phosphorylated by the kinase Aurora A, which is essential for normal MT flux on the metaphase spindle. NuSAP knockout in cells leads to the improper increase of Kif2A localization onto spindle poles while reduces Eg5 localization on spindle microtubules. Moreover, NuSAP phosphorylation by Aurora A at mitosis enhances its affinity with Kif2A on spindle microtubules, thus preventing superabundant concentration of Kif2A onto the spindle poles and maintaining the proper metaphase microtubule flux. Taken together, our study reveals a novel mechanism of how NuSAP regulates mitotic spindle assembly, structural dynamics, and metaphase spindle length control for accurate chromosome congression via affecting MT flux and Kif2A localization.

## Results

### NuSAP deletion results in short spindle formation, whereas its overexpression makes the spindle longer

To understand how cells control spindle assembly and their chromosome congression, we explored the roles of NuSAP in spindle dynamics and function. First, we established NuSAP knockout cell lines with CRISPR-Cas9 in HeLa cell lines. Immunofluorescence microscopy (IFM) and western blot analysis showed that we successfully knocked out the expression of NuSAP in HeLa cells (*Figure 1A-B*). Through IFM, we observed that NuSAP knockout showed a remarkable effect rin generating short bipolar metaphase spindles (*Figure 1A*). The mean distance between spindle poles, as marked by γ-tubulin signals, was reduced to ~10.07 μm compared with the normal length of ~11.23 μm (*Figure 1C*). Under conditions that overexpress NuSAP in cells, the metaphase spindle length was significantly longer (~14.43 μm) than in control cells (~11.57 μm) (*Figure 1D-E*). Notably, overexpression of NuSAP also induced unusual curved MT filaments, extensive bundling of the MTs and deficient spindle morphology (*Figure 1D*), as reported before (Raemaekers et al., 2003). These results suggest that proper amount of NuSAP, not more or less, is essential for a precise metaphase spindle assembly like through participating in the metaphase spindle length control.

**Figure 1.**
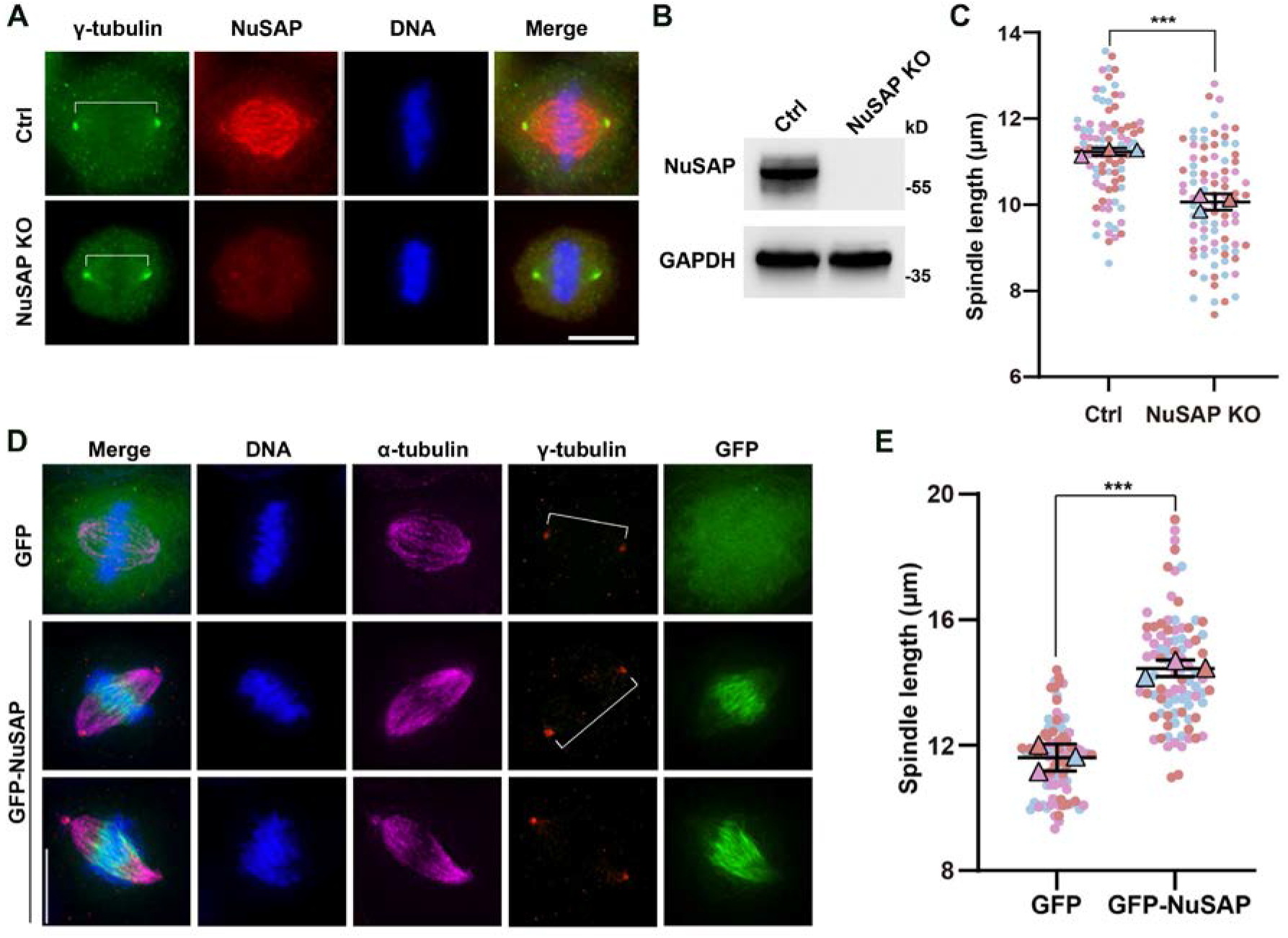
NuSAP deletion results in short bipolar spindle formation, whereas its overexpression makes the spindle longer. (A) Control and NuSAP-knockout HeLa cells were stained for γ-tubulin (green), NuSAP (red), and DNA (blue). (B) NuSAP knockout efficiency of HeLa cells was analyzed by Western blotting. (C) Pole-to-pole distances (indicated by white broken lines) were quantified by γ-tubulin immunofluorescence in control or NuSAP-knockout metaphase cells. Error bars indicated SD. Three independent replicates of 30 cells per replicate were quantified. Unpaired two-tailed t test: P = 0.0006. (D) HeLa cells expressing GFP or GFP-NuSAP were stained for γ-tubulin (red), α-tubulin (magenta) and DNA (blue). (E) Pole-to-pole distances (indicated by white broken lines) were measured by γ-tubulin immunofluorescence in metaphase cells Error bars indicated SD. Three independent replicates of 30 cells per replicate were quantified. Unpaired two-tailed t test: P = 0.0006. Figure 1-source data 1 Raw Microsoft excel file used for analysis of graphs in Figure 1C and E. Figure 1-source data 2 Labeled uncropped western blot images and raw western blot images in Figure 1B.

### NuSAP depletion accelerates metaphase spindle MT flux

MT poleward flux can serve as an important factor in spindle dynamics and controls spindle elongation (Buster et al., 2007; Cimini, Wan, Hirel, & Salmon, 2006; Laycock, Savoian, & Glover, 2006; Maffini et al., 2009; Steblyanko et al., 2020). Based on our observations above, we tested whether NuSAP knockout affects the poleward translocation/flux of tubulin dimers along the spindle MTs. We transfected control and NuSAP-knockout HeLa cells with photoactivatable (PA) GFP-tagged α-tubulin. Then we used pulse flash of 405 nm laser to activate a rectangular region of PAGFP-α-tubulin in the proximity of chromosomes while time-lapse images were captured every 10 s to measure the velocity at which the activated fluorescent signals approached the pole (*Figure 2A-B, Video 1 and Video 2*). In control metaphase cells, the fluorescent signals approached the pole with a mean velocity of 0.30 μm/min. In contrast, after NuSAP knockout, the velocity of MT flux was significantly increased to 0.44 μm/min (*Figure 2C-F*). Thus, these results suggest that NuSAP deletion acc MT flux of metaphase spindle. However, previous study shows that MT-flux rate and spindle length showed a strong positive correlation overall. Accordingly, NuSAP deletion may affect other spindle length regulators to counteract spindle elongation caused by MT flux rate increase.

**Figure 2.**
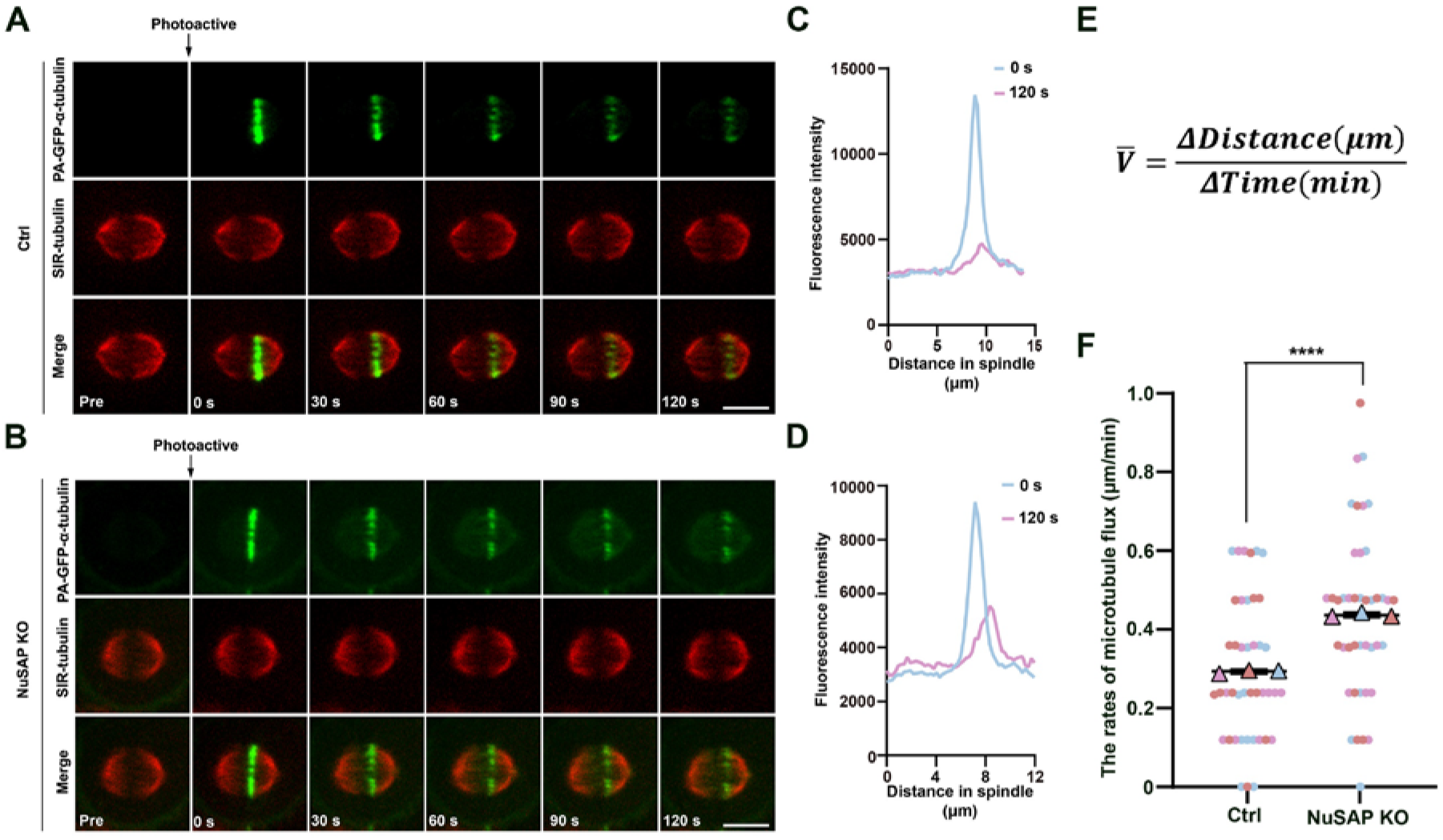
NuSAP depletion slows down spindle MT flux. (A and B) NuSAP-knockout HeLa cells were transfected with photoactivatable GFP-tagged α-tubulin (PAGFP-α-tubulin). Microtubules were stained with SiR-tubulin. GFP signal in a rectangular region near the MT-plus ends was activated (time point 0, arrows) and tracked every 10 s. Representative time-course images are shown. (C and D) The fluorescence intensity profiles at time point 0 s and 120 s in (A and B) were shown respectively. (E) The mean velocity was defined by the ratio of the distance that the PAGFP signal travels to 2 min it takes. Δ Distance: the distance between the two points with the strongest PAGFP fluorescence intensity at 0 s and 120 s. Δ Time: 2 min. (F) The rates of MT flux shown in (A and B) were measured in control or NuSAP-knockout cells. Error bars indicated SD. Three independent replicates of 14 cells per replicate were quantified. Unpaired two-tailed t test: P < 0.0001. ***, P < 0.001; ****, P < 0.0001. Scale bars, 10 μm. Figure 2-source data 1 Raw Microsoft excel file used for analysis of graphs in Figure 2C, D and F.

Proteins that influence MT polymer dynamics can cause the numbers and average length of spindle MTs to be anticorrelated (Goshima & Kimura, 2010). In addition to the centrosome-based MT nucleation, MTs were also shown to be generated within the spindle independently of centrosomes by MT nucleation adjacent to the chromosomes (W. Fu et al., 2013; Tulu, Rusan, & Wadsworth, 2003) or MT-based MT nucleation process mediated by γ-tubulin ring complex (γ-TuRC) and its spindle-localizing factor (Goshima, Mayer, Zhang, Stuurman, & Vale, 2008; Petry, Groen, Ishihara, Mitchison, & Vale, 2013; Song et al., 2018). These acentrosomal small asters contribute to the spindle assembly through interacting with each other, binding and sorting into the big centrosomal MT asters (W. Fu et al., 2013). Observations that disruption of this process leads to the formation of unusually long metaphase spindles support the hypothesis that acentrosomal MT nucleation within spindles is important for spindle length control (Goshima et al., 2008). To verify whether NuSAP regulates the spindle length through mediating the acentrosomal MT nucleation, we treated HeLa cells with 500 ng/mL nocodazole for 2 h to disassemble overall MTs followed by releasing these cells into fresh medium containing a low concentration of nocodazole (15 ng/mL) to induce the acentrosomal MT nucleation (W. Fu et al., 2013). Through immunostaining of the fixed cells, we observed the acentrosomal microtubule assembly, and the results showed that whereas the centrosome-based MT nucleation was partially inhibited in the presence of this low concentration of nocodazole, the acentrosomal MT nucleation was induced, and these nucleated MTs rapidly assembled into many small acentrosomal asters (*Figure 2-figure supplement 1A*). The analysis results showed that the number of small MT asters in NuSAP knockdown cells was no different from that in control cells (*Figure 2-figure supplement 1B-C*). Thus, the results indicate that NuSAP controlling the metaphase spindle length is not via mediating acentrosomal MT nucleation.

### NuSAP depletion or knockdown also results in kinetochore-MT attachment failure

The proper dynamic turnover of spindle MTs is essential for a stable connection between kinetochores and MTs (Cheeseman & Desai, 2008). We have shown that NuSAP depletion leads to spindle MTs less stable and faster poleward movement. Here, we asked whether NuSAP contributes to the kinetochore-MT connection during mitosis. First, mitotic HeLa cells transfected with control or NuSAP siRNA, treated with MG132 to achieve fully assembled bipolar spindles, and placed on ice for 10 min before fixation. With IFM, we observed that the intact end-on attached K-fibers were stably preserved in normal control cells under cold treatment; in contrast, few intact K-fibers were left in NuSAP knockdown cells (*Figure 3A-B*), suggesting that NuSAP knockdown abolished the stable connection between kinetochores and MTs during mitosis. Consistently, we uncovered that the inter-kinetochore distance of chromosomes aligned at the metaphase plate in NuSAP knockdown cells (0.91 ± 0.01 μm) was substantially shorter than that in control metaphase cells (1.17 ± 0.02 μm), indicating a lack of tension between a pair of kinetochores and that NuSAP was also required for maintenance of proper inter-kinetochore tension (*Figure 3C-D*). To understand the underlying the mechanism, we stained the cells with a specific antibody against the spindle checkpoint protein BubR1. The results showed that BubR1 was maintained at the kinetochores in NuSAP knockdown cells at metaphase (*Figure 3E*),suggesting that a proper stable connection between kinetochores and spindle MTs was not well established. Taking these findings together, we conclude that NuSAP-regulated MT dynamics contributes to the kinetochore-MT attachment during mitotic spindle assembly, structural dynamics, and chromosome congression.

**Figure 3.**
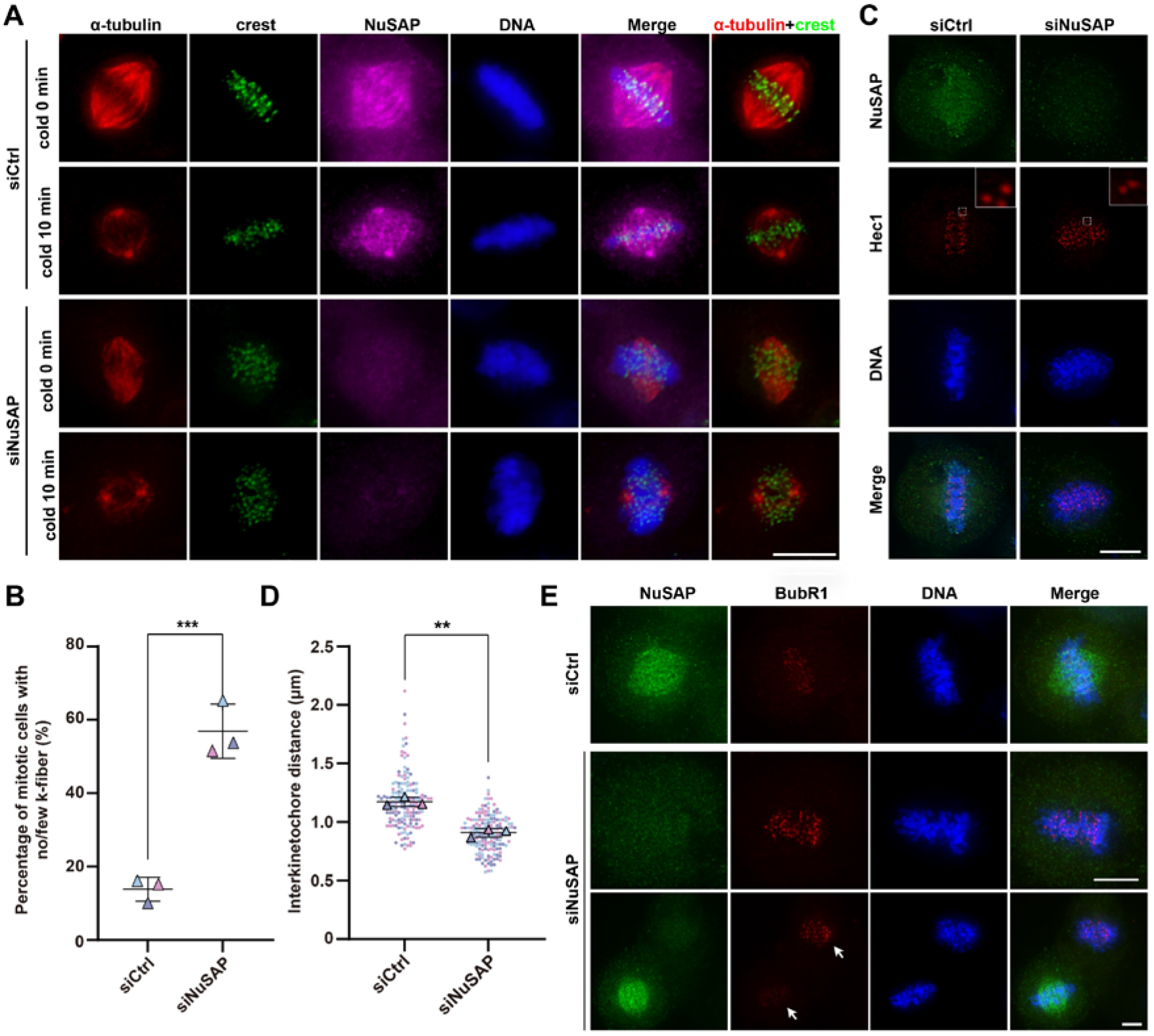
NuSAP depletion or knockdown also results in kinetochore-MT attachment failure. (A) HeLa cells transfected with control or NuSAP siRNA were placed on ice for 10 min before fixation. Then the cells are processed for IFM using anti-α-tubulin (red), anti-crest (green) and anti-NuSAP (magenta) antibodies. Note that NuSAP knockdown reduced the k-fibers, resulting in significant chromosome misalignment. (B) Quantification of the percentage of mitotic cells with no/few k-fibers. Error bar indicated SD. Unpaired two-tailed t test. P=0.0008. 303 cells and 219 cells were counted in control cells and NuSAP knockdown cells from three independent experiments. (C) HeLa cells with NuSAP knockdown were immunostained with anti-NuSAP (green) and anti-Hec1 (red) antibodies. The box areas were zoomed at the top right. (D) Measurement of interkinetochore (labeled by Hec1) distances. Error bars indicated SD. P=0.0011. Three independent replicates of 53 pairs of kinetochores per replicate were quantified. Unpaired two-tailed t test. (E) Knockdown of NuSAP by siRNA in HeLa cells led to activation of the spindle assembly checkpoint. HeLa cells with siRNA knockdown of NuSAP were stained with anti-NuSAP (green) and anti-BubR1 (red) antibodies. Note that NuSAP knockdown led to chromosome misalignment and BubR1-positive staining (arrow). The DNA was stained by DAPI (blue) in (A, C and E). **, P < 0.01; ***, P < 0.001. Scale bars, 10 μm. Figure 3-source data 1 Raw Microsoft excel file used for analysis of graphs in Figure 3B and D.

### NuSAP is phosphorylated in mitosis by Aurora A

To address how the function of NuSAP is regulated, we examined the levels of the NuSAP protein in different phases of the cell cycle. HeLa cells were arrested at the G1/S boundary by a double-thymidine treatment and then released into fresh medium and collected samples at different time points. Western blot analysis showed NuSAP is a cell cycle-depended protein with a high level of expression in the G2/M phase, followed by a sharp decline in the G1 phase. Moreover, a portion of NuSAP proteins exhibited a motility shift on SDS-PAGE when cells enter mitosis (indicated by Histone H3pS10 antibody) (*Figure 4A*), which suggested that NuSAP might be posttranslational modified during mitosis progression. The mobility change was abolished when the cell mitotic lysates were pretreated with λ-phosphatase (*Figure 4B*), suggesting that NuSAP was phosphorylated during mitosis.

**Figure 4.**
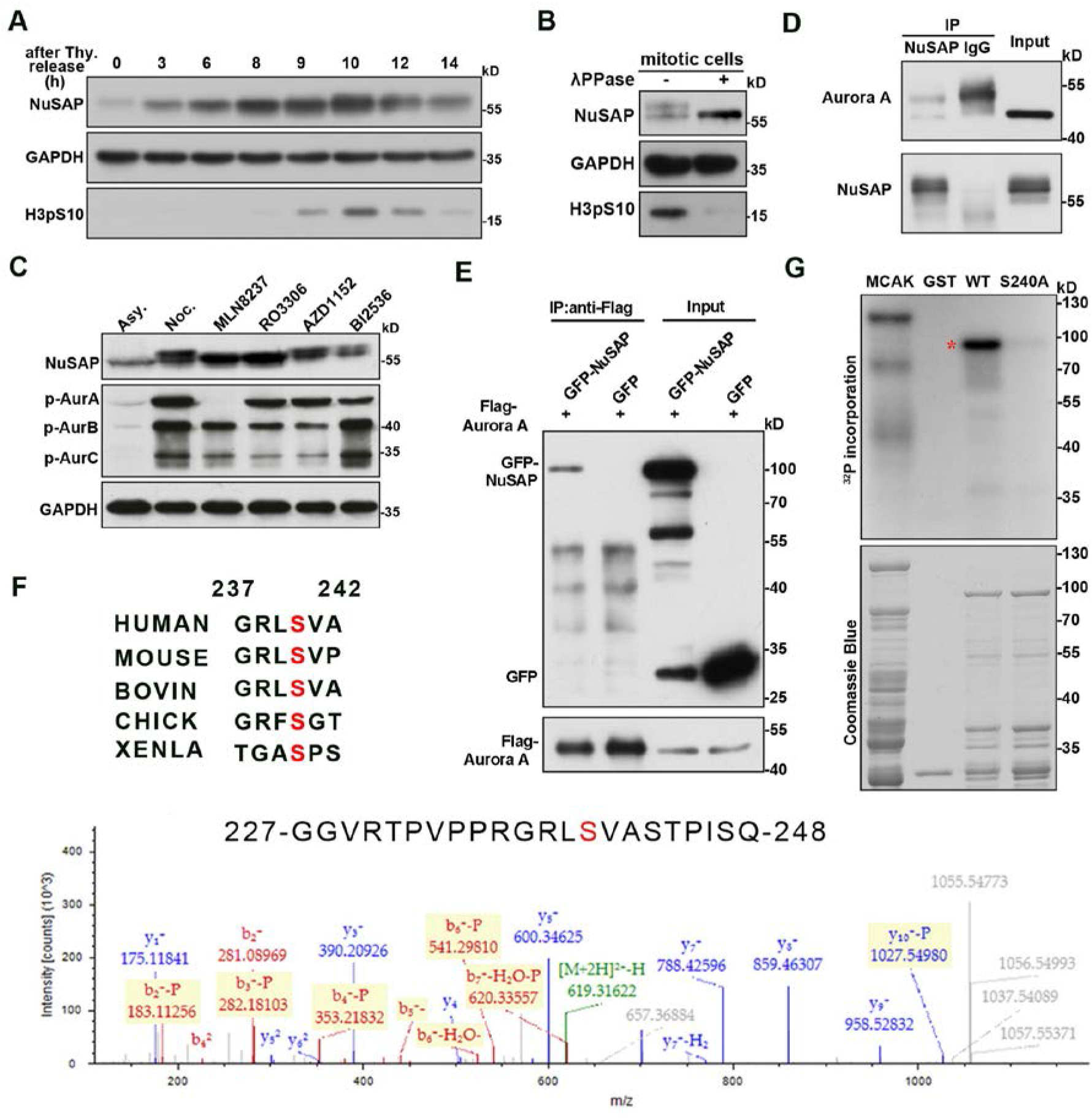
NuSAP is phosphorylated at S240 by Aurora A in mitosis. (A) HeLa cells were blocked at G1/S by double thymidine (Thy.) treatment and then released into fresh medium and harvested at the indicated time point. (B) Mitotic HeLa cell lysates were treated with λ-phosphatase. Samples were analyzed on SDS-PAGE and immunoblotted with the indicated antibodies. (C) HeLa cells were arrested with nocodazole for 17 h and then treated with selected kinase inhibitors (0.25 μM MLN8237 for 30 min, 9 μM RO3306 for 15 min, 0.1 μM AZD1152 for 30 min, 0.1 μM BI2536 for 30 min, respectively.) Asy., asynchronized cell samples. Samples were analyzed on SDS-PAGE and immunoblotted with the indicated antibodies. (D) Mitotic HeLa cells were subjected to an immunoprecipitation (IP) assay with nonspecific IgG or anti-NuSAP anti-body followed by Western blotting. (E) Flag-Aurora A was co-transfected with GFP or GFP-NuSAP into HEK293T cells. Lysates of transfected 293T cells were subjected to IP assay with antiFlag antibody followed by western blotting. (F) GST-tagged NuSAP protein was incubated with Aurora A kinase in 30°C for 30 min. Samples were processed for MS analysis. Ser-240 site was found to be phosphorylated, which is consistent with the Aurora A phosphorylation consensus motif. Multiple sequence alignment in top panel was performed using Uniprot. Red amino acids were conserved. (G) GST-tagged NuSAP proteins with/without point mutation were subjected to Aurora A kinase assay in vitro followed by autoradiography (Markopoulos et al.). GST-tagged MACK was used as positive control. Coomassie Blue (right) staining shows the loading of the GST-tagged NuSAP proteins in the reactions. Phosphorylated recombinant NuSAP is indicated by an asterisk. Figure 4-source data 1 Raw Microsoft excel file for mass spectrum data in Figure F (left panel). Figure 4-source data 2 Labeled uncropped western blot images, raw western blot images, Coomassie stained gel images and full autoradiography images in Figure 4A-E, and G.

We then screened the possible mitotic kinases responsible for NuSAP phosphorylation. Through treating mitotic HeLa cells with several mitotic kinase inhibitors, we revealed that treatment with the specific inhibitor MLN8237 for kinase Aurora A and RO3306 for kinase Cdk1 significantly reduced the higher band of NuSAP; in contrast, Aurora B kinase inhibition by AZD1152 or Plk1 kinase inhibition by BI2536 did not down-shift the NuSAP band (*Figure 4C*). A previous report has shown that Cdk1 is responsible for NuSAP phosphorylation at T300 and T338 sites during mitosis, whereas this phosphorylation inhibits NuSAP binding with microtubules (Chou et al., 2011). Thus, we focused on the functions of NuSAP phosphorylation by Aurora A. First, we wanted to know whether NuSAP and Aurora A could bind with each other *in vivo*. We performed an immunoprecipitation (IP) assay using anti-NuSAP antibody and found that endogenous Aurora A could be immunoprecipitated from mitotic lysates (*Figure 4D*). In a parallel experiment, we co-expressed Flag-Aurora A with GFP or GFP-NuSAP in HEK293T cells followed by IP assay, the results also show that NuSAP interacted with Aurora A (*Figure 4E*). To identify the phosphorylation site(s) of Aurora A on NuSAP, we expressed and purified GST-tagged NuSAP from *E.coli* and incubated the purified proteins with Aurora A kinase *in vitro* treated by DMSO or MLN8237, then prepared the samples for mass spectrum (MS) analysis. The results revealed that Ser-240, which is consistent with the consensus sequence R-X-pS/T-L/V of Aurora A (Ohashi et al., 2006), was phosphorylated (*Figure 4F, bottom panel*). Besides, the site is conserved in different species, from human to *Xenopus* (*Figure 4F, top panel*), although it is not in an Aurora A consensus phosphorylation site in *Xenopus*. To further verify this phosphorylation, we constructed and purified NuSAP mutant protein NuSAP-S240A and performed an *in vitro* kinase assay. The results showed that NuSAP-WT was phosphorylated by Aurora A kinase while the mutant NuSAP-S240A could hardly be phosphorylated (*Figure 4G*), indicating that S240 contributed to the proper phosphorylation. This result was also consistent with previous reports (Sardon et al., 2010), whereas the related functions are not clear..

### Phosphorylation status of NuSAP by Aurora A affects mitotic spindle dynamics via regulating spindle MT flux

To determine the biological importance of NuSAP phosphorylation by Aurora A, we performed the rescue assay in NuSAP knockout HeLa cells using GFP-NuSAP-WT, GFP-NuSAP-S240A or GFP-NuSAP-S240D (*Figure 5A-B*). Through immunostaining for α-tubulin in NuSAP-knockout and-rescue cells, we tested how S240 phosphorylation of NuSAP regulates mitotic spindle dynamics. The results showed that GFP-tagged NuSAP-WT and NuSAP-S240D almost fully rescued metaphase short spindles resulting from NuSAP knockout, which even caused spindle elongation beyond its normal length due to the overexpression, whereas NuSAP-S240A was unable to rescue this defect (*Figure 5A-B*). Furthermore, we carried out live-cell imaging in NuSAP-knockdown HeLa cells while expressing different NuSAP mutants and RFP-H2B, and revealed delayed chromosome biorientation and mitosis progression in cells transfected with GFP-NuSAP-S240A. The mean time from NEBD to proper chromosome alignment in cells expressing NuSAP-WT or NuSAP-S240D was 61.40 ± 3.72 min, and 96.70 ± 7.76 min, respectively, while this time was significantly prolonged to 203.4 ± 18.98 min in cells expressing NuSAP-S240A (*Figure 5-supplement 1A-B, Video 3–5*). Taken together, these results demonstrate that the phosphorylation of NuSAP at Ser-240 by Aurora A regulates the spindle assembly, dynamics, and chromosome congression for normal mitotic progression.

**Figure 5.**
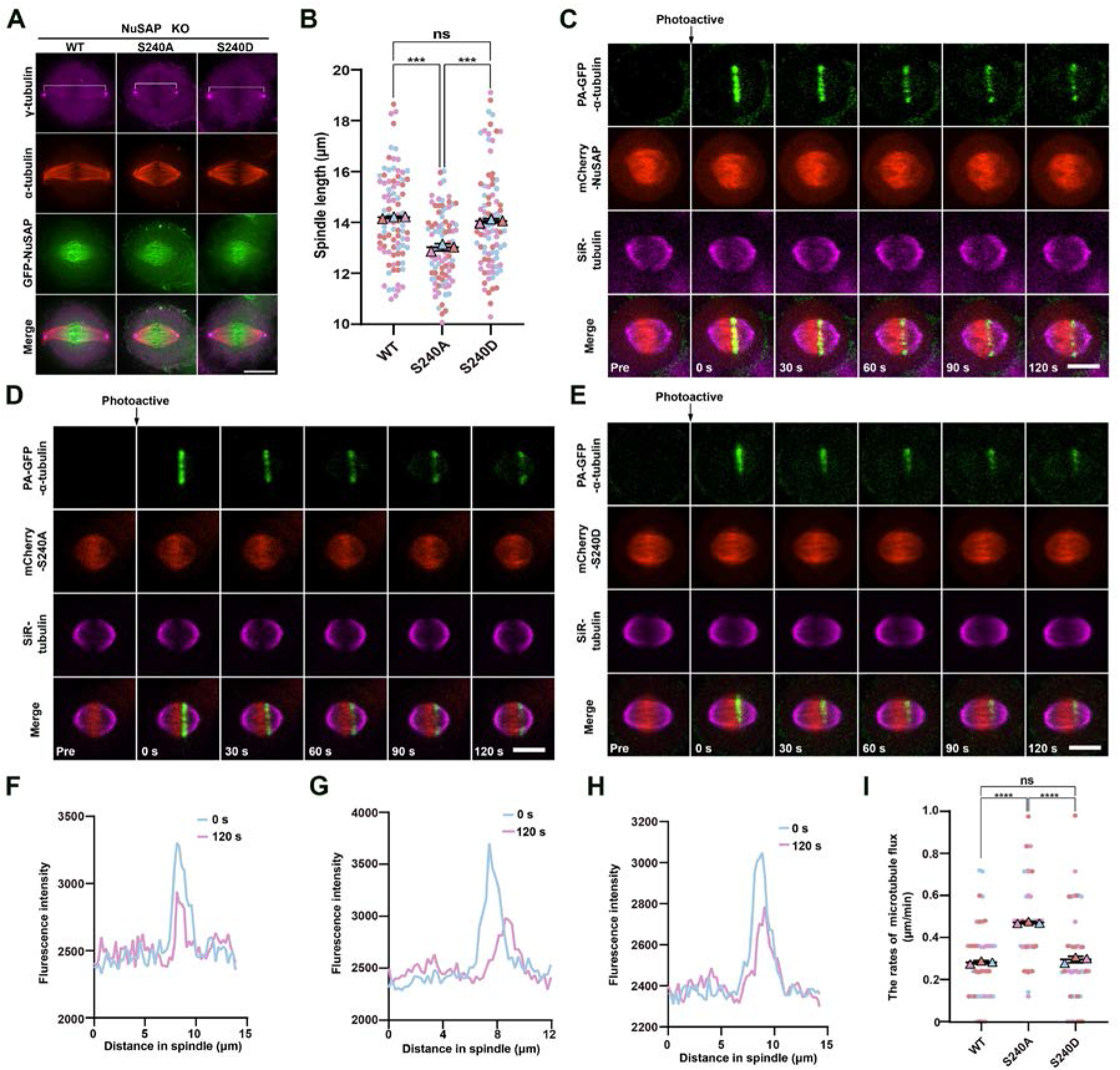
Phosphorylation status of NuSAP by Aurora A affects mitotic spindle dynamics via regulating spindle MT flux. (A) NuSAP-knockout HeLa cells were transfected with GFP-tagged NuSAP-WT, -S240A, or -S240D for 48 h followed by immunofluorescence labeling using anti-α-tubulin antibody and anti-γ-tubulin antibody. (B) Quantification of spindle length in (A). Error bars indicated SD. Three independent replicates of 30 cells per replicate were quantified. Unpaired two-tailed t test: P = 0.0002 for GFP-NuSAP/GFP-S240A; P = 0.0004 for GFP-S240A/GFP-S240D; P = 0.0550 for GFP-NuSAP/GFP-S240D. ns, not significant; ***, P < 0.001. Scale bars, 10 μm. (C-E) NuSAP-knockout HeLa cells were transfected with photoactivatable GFP-tagged α-tubulin (PAGFP-α-tubulin) and mCherry-tagged NuSAP-WT, -S240A, or −240D followed by live-cell imaging. GFP signal in a rectangular region near the MT plus ends was activated (time point 0, arrows) and tracked every 10s. Representative time-course images are shown. (F-H) The fluorescence intensity profiles at time point 0 s and 120 s in (C-E) were shown respectively. (I) The rates of MT flux shown in (C-E) were measured (as described in the legend for Fig. 2F). Error bars indicated SD. Three independent replicates of 13 or 14 cells per replicate were quantified. Unpaired two-tailed t test: P < 0.0001 for GFP-NuSAP/GFP-S240A; P < 0.0001 for GFP-S240A/GFP-S240D; P = 0.2511 for GFP-NuSAP/GFP-S240D. ns, not significant; ****, P < 0.0001. Scale bars, 10 μm. Figure 5-source data 1 Raw Microsoft excel file used for analysis of graphs in Figure 5B, F-I.

As we have observed that the NuSAP knockout lead to the increased rate of MT flux, we asked whether the phosphorylation of NuSAP also regulates this process. We cotransfected NuSAP-knockout cells with PAGFP-α-tubulin and different mCherry-tagged NuSAP constructs and performed the photoactivation experiment as previous (*Figure 5C-E, Video 6–8*). Through measuring the velocity of fluorescent signal approached the pole, we found that in NuSAP knockout metaphase cells, the mean velocity of MT flux is 0.28 μm/min when expressing mCherry-NuSAP-WT, and is 0.30 μm/min when expressing mCherry-NuSAP-S240D, whereas the rate of poleward flux was significantly increased to 0.47 μm/min when expressing NuSAP-S240A (*Figure 5F-I*). Hence, we concluded that the phospho-null NuSAP mutant could not rescue the defects in MT flux caused by NuSAP knockout. Together, these results show that phosphorylation of NuSAP Ser-240 maintains normal spindle microtubule flux, while phospho-null NuSAP accelerates the poleward movement of MTs.

### NuSAP negatively regulates the concentration of Kif2A on spindle poles and promotes the localization of Eg5 on spindle MTs

To investigate how NuSAP might affect spindle length, we next identified the cellular targets of NuSAP in mitosis. Firstly, we tested the interaction between NuSAP and several established contributors to MT flux and spindle length control, including Kif2A, MCAK, Eg5, and KifC1. Kif2A and MCAK were able to depolymerize microtubules on spindle poles and kinetochores respectively and so to regulate spindle length (Ganem & Compton, 2004). Especially, it has been reported that NuSAP regulates the dynamics of kinetochore microtubules by attenuating MCAK depolymerization activity (C. Li et al., 2016). Eg5 contributes spindle MT flux by mediating sliding of antiparallel interpolar MTs, and the changes in spindle length caused by knockdown or overexpression of KifC1 are relatively similar with that of NuSAP (Brust-Mascher et al., 2009; Cai, Weaver, Ems-McClung, & Walczak, 2009; Miyamoto et al., 2004). Surprisingly, the IP assay showed that NuSAP could interact with all the four protein regulators mentioned above (*Figure 6-supplement 3A*). Next, we explored whether they were regulated by NuSAP. We analyzed the localization of these regulators after knockout or overexpression of NuSAP. IFM statistics showed that NuSAP had no effect on the relative fluorescence intensity of KifC1 on spindle MTs and that of MCAK at microtubule plus ends (*Figure 6-supplement 3B-E*). In contrast, Kif2A fluorescence intensity on spindle poles was significantly enhanced due to NuSAP knockout and reduced because of NuSAP overexpression (*Figure 6A-D*). And on spindle MTs localized Eg5 was significantly reduced in NuSAP-knockout cells compared with the control, whereas it was also reduced by NuSAP overexpression, which was probably caused by spindle elongation (*Figure 6E-H*). To confirm the interaction between NuSAP and the two target proteins, we performed an IP assay using anti-NuSAP antibodies and pulled down endogenous Eg5 or Kif2A from mitotic HeLa lysates. Indeed, the results showed that endogenous NuSAP interacted with endogenous Eg5 and Kif2A (*Figure 6I-J*). Though western blot analysis, we showed that the protein levels of Eg5 and Kif2A were not affected by NuSAP knockdown (*Figure 6K*). These results indicate that NuSAP just influenced the subcellular localization of Eg5 and Kif2A but not their expression. Furthermore, we co-depleted Kif2A and NuSAP in HeLa cells and measured the rates of microtubule flux. We found that the increased rate of microtubule flux caused by NuSAP knockout was rescued by Kif2A depletion, indicating that NuSAP regulated MT flux mostly through Kif2A (*Figure 6L*). Collectively, these results indicate that NuSAP negatively regulated the concentration of Kif2A on spindle poles and promoted the localization of Eg5 on spindle MTs, so Eg5 and Kif2A might be the main downstream targets of NuSAP.

**Figure 6.**
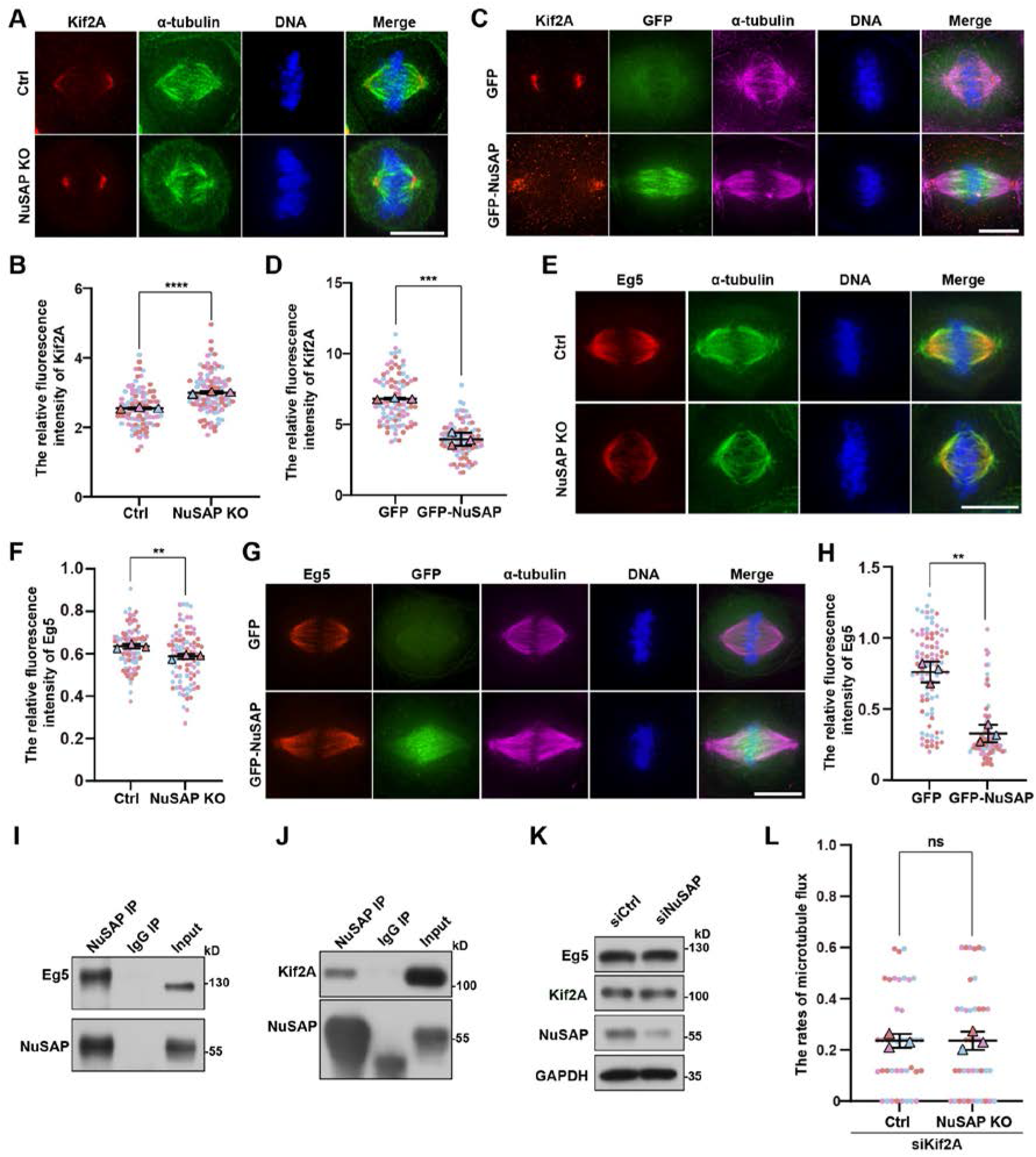
NuSAP negatively regulates the concentration of Kif2A on spindle poles and promotes the localization of Eg5 on spindle MTs. (A and E) Control and NuSAP-knockout HeLa cells were stained with Kif2A (red), α-tubulin (green) in (A), and stained with Eg5 (red), α-tubulin (green) in (E). DNA was stained with DAPI (blue). (B and F) Quantification of relative Kif2A (B) and Eg5 (F) fluorescence intensity on spindle in (A) and (E), respectively. Error bars indicated SD. Three independent replicates of 30 cells per replicate were quantified. Unpaired two-tailed t test: P < 0.0001 in (B); P = 0.0080 in (F). (C and G) Mitotic HeLa cells were transfected with GFP or GFP-NuSAP and stained with Kif2A (red), α-tubulin (purple) in (C), and stained with Eg5 (red), α-tubulin (purple) in (G). DNA was stained with DAPI (blue). (D and H) Quantification of relative Kif2A (D) and Eg5 (H) fluorescence intensity on spindle in (C) and (G), respectively. Error bars indicated SD. Three independent replicates of 30 cells per replicate were quantified. Unpaired two-tailed t test: P = 0.0004 in (D); P = 0.0014 in (H). (I and J) Mitotic HeLa cells were subjected to an IP assay using nonspecific IgG or anti-NuSAP antibody followed by western blot analysis with anti-NuSAP, anti-Eg5 (I) and anti-Kif2A (J) antibodies. (K) Test the protein level of Kif2A and Eg5 after NuSAP knockdown in HeLa cells. (L) The mean velocity of poleward spindle microtubule flux in Kif2A-depleted and NuSAP/Kif2A-co-depleted cells. The mean velocity was defined by the ratio of the distance that the PAGFP signal travels to the time it takes. Error bars indicated SD. Three independent replicates of 13 or 14 cells per replicate were quantified. Unpaired two-tailed t test: P = 0.9884. ns, not significant; **, P < 0.01; ***, P < 0.001; ****, P < 0.0001. Scale bars, 10 μm. Figure 6-source data 1 Raw Microsoft excel file used for analysis of graphs in Figure 6B, D, F, H, and L. Figure 6-source data 2 Labeled uncropped western blot images and raw western blot images in Figure 6I-K.

### NuSAP controls spindle assembly, dynamics, and metaphase spindle length by negatively regulating Kif2A on spindle poles in an Aurora A-dependent manner

To further test whether phosphorylation of NuSAP at Ser-240 affects these interactions, we co-transfected cells with GFP-Kif2A and different mCherry-tagged NuSAP constructs followed by co-IP assay in mitosis cells. The results showed that Kif2A has stronger affinity to phosphorylated NuSAP, including WT and S240D (*Figure 7A-B*). In the same way, we found that Eg5 bound to NuSAP-WT, -S240A, and -S240D with similar affinity (*Figure 7C-D*). These data show that the interaction between NuSAP and the MT depolymerase Kif2A is Aurora A phosphorylation-dependent and that abolishing NuSAP phosphorylation reduces its affinity to Kif2A. Next, through immunostaining for Kif2A in NuSAP knockout-and-rescue cells, we further tested that how phosphorylation of NuSAP at Ser-240 influences Kif2A function in regulating metaphase MT flux and spindle assembly. To fully establish the bipolar spindle, we treated synchronized mitotic cells with 10 μM MG132 for 1 h. The results showed that more Kif2A localized on spindle poles in cells expressing NuSAP-S240A compared with that in cells expressing NuSAP-WT and - S240D (*Figure 7E-F*). As reported, Kif2A could depolymerize MTs at their minus ends (Desai, Verma, Mitchison, & Walczak, 1999; Ganem, Upton, & Compton, 2005). Consequently, increased concentration of Kif2A on spindle MTs, which caused by NuSAP knockout or NuSAP-240A overexpression would accelerate the MT flux on spindle. Taken together, these results indicate that NuSAP controls the spindle assembly, structural dynamics, and metaphase length by negatively regulating Kif2A on spindle poles in an Aurora A-dependent manner.

**Figure 7.**
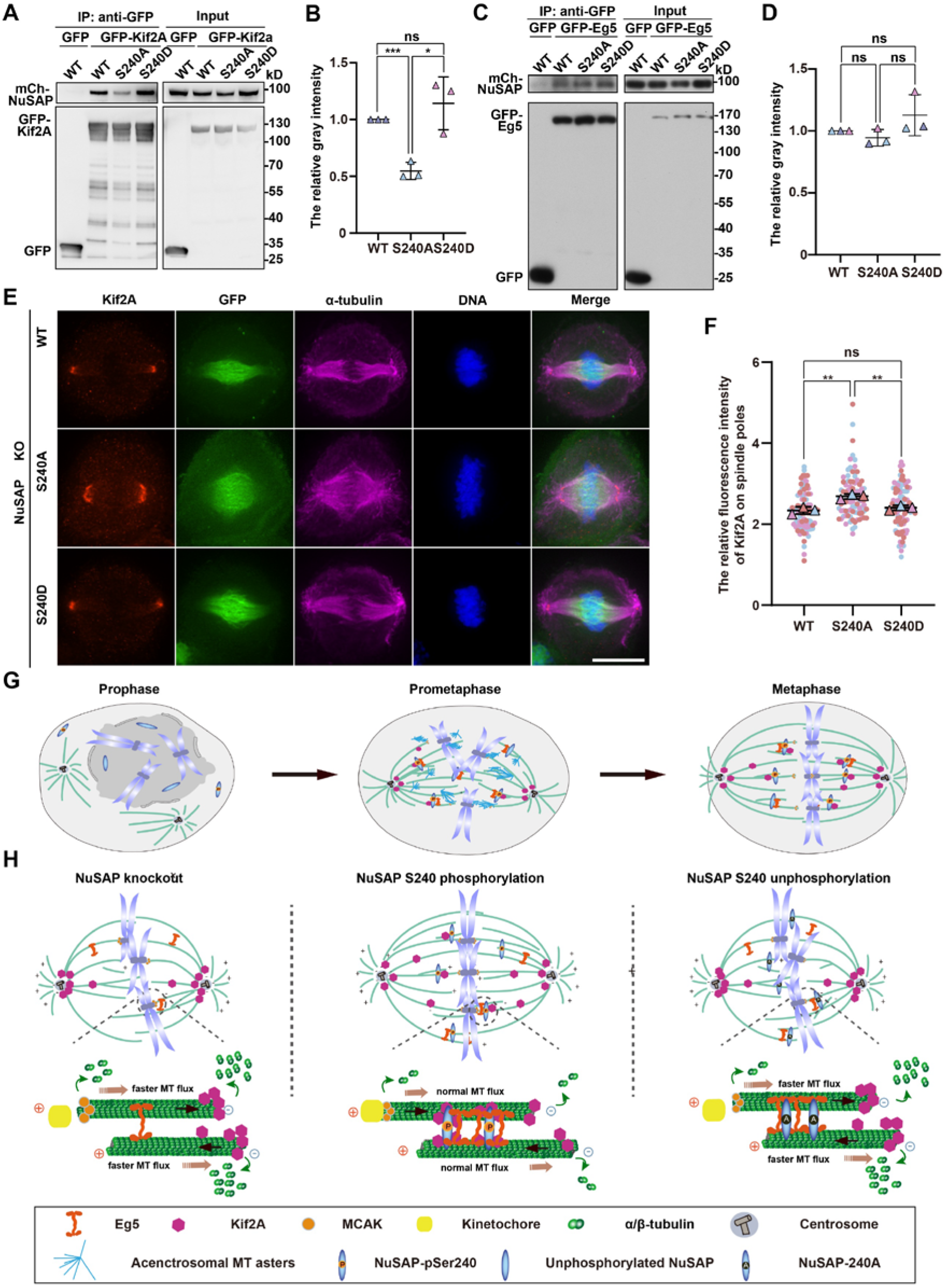
Ser-240 phosphorylation of NuSAP promotes its interaction with Kif2A and reduces localization of Kif2A on spindle poles. (A and C) Mitotic HEK293T cells co-transfected with the indicated GFP-tagged proteins and mCherry-tagged NuSAP constructs were immunoprecipitated with GFP-Trap beads. NuSAP-S240A binds significantly less Kif2A than -WT and -S240D, while NuSAP-WT, -S240A, and -S240D show similar affinity to Eg5. (B and D) The protein-protein interaction discrepancy was quantified by ImageJ. The relative interaction between mCherry-tagged NuSAP constructs and GFP-Kif2A (B) or GFP-Eg5 (D) was defined by the ratio of the amount of interacting mCherry-NuSAP proteins to that of immunoprecipitated GFP-Kif2A or GFP-Eg5. The value of the WT group was set as 1.0, and the other mutant groups were calculated. Error bars indicated SD. Three independent replicates were quantified. Unpaired two-tailed t test: P = 0.0005 for WT/ S240A, P = 0.0135 for S240A/S240D, P = 0.3489 for WT/ S240D in (B); P = 0.2274 for WT/S240A, P = 0.1531 for S240A/S240D, P = 2562 for WT/S240D in (D). (E) NuSAP-knockout HeLa cells were transfected with GFP-tagged NuSAP-WT, -S240A, or -S240D followed by immunofluorescence labeling using anti-Kif2A (red) and anti-α-tubulin (purple) antibodies. DNA was stained with DAPI (blue). (F) Quantification of relative Kif2A intensity on spindle poles in (E). Error bars indicated SD. Three independent replicates of 13 or 14 cells per replicate were quantified. Unpaired two-tailed t test: P = 0.0049 for WT/S240A; P = 0.0051 for S240A/S240D; P = 0.3347 for WT/ S240D. ns, not significant; *, p < 0.05; **, P < 0.01; ***, P < 0.001. Scale bars, 10 μm. (G and H) A working Model showing NuSAP functions in regulating the mitotic spindle dynamics and length control. (G). Along with the nuclear envelope broken down during mitotic entry, MTs asters nucleate around the chromosomes and the centrosomes to form small acentrosomal and large centrosomal asters. These oriented MTs are bundled then and sorted into the large centrosomal MT asters to form a bipolar spindle structure by molecular motors and MT-associated proteins till to well-organized metaphase spindle formation with a dynamically constant length, in which NUSAP plays a crucial role. (H). NuSAP is mainly localized on MTs near chromosomes and is phosphorylated by the mitotic kinase Aurora A on S240. As a MAP, NuSAP regulates the spindle dynamics to maintain metaphase spindle length through coordinating MT depolymerization and MT-MT sliding. On one hand, NuSAP interacts with Eg5 on the adjacent MTs, which allows Eg5 to move along the MTs and to mediate the sliding of antiparallel interpolar MTs (shown by the black arrow), which increases spindle length. On the other hand, S240 phosphorylated NuSAP interacts with more Kif2A on the spindle body and then reduces the amount of Kif2A on spindle poles, thus restraining the excess accumulation of Kif2A at the minus ends of MTs to ensure the proper microtubule depolymerization. At the plus end, NuSAP attenuates MCAK depolymerization activity to stabilize kinetochore microtubules. NuSAP regulates microtubule flux (shown by the orange arrow) and controls metaphase spindle length by coordinating with Eg5, Kif2A, and MCAK. In experimental condition, NuSAP knockout results in a reduction of Eg5 localization on spindle microtubules, excess accumulation of Kif2A on spindle poles, and high MCAK depolymerization activity, which leads to faster microtubule flux, kinetochore-microtubule attachment failure, and thus to formation of a shorter metaphase spindle. Alternatively, overexpression of NuSAP-S240A, a loss-of-function mutant that interacts with less Kif2A on spindle body, also causes faster microtubule flux and shorter metaphase spindle formation. Figure 7-source data 1 Raw Microsoft excel file used for analysis of graphs in Figure 7B, D, and F. Figure 7-source data 2 Labeled uncropped western blot images and raw western blot images in Figure 7A and C.

## Discussion

Based on our present work and previous reports (J. Fu et al., 2015; Ma et al., 2010; Zhuo et al., 2015), we propose a working model to illustrate the underlying mechanisms on the regulation of spindle assembly, structural dynamics, and metaphase spindle length control (*Figure 7G-H*). After nuclear envelope breakdown, MTs nucleate around chromosomes and centrosomes and grow with dynamic instability. These oriented MTs are bundled with each other and gradually sorted into a bipolar spindle by molecular motors and non-motor proteins. During this process, NuSAP, serving as a MAP, not only promotes kinetochore-MT attachment for K-fiber formation and chromosome congression and coordinates MT depolymerization and MT-MT sliding, but also interacts with Eg5 on spindle MTs, which allows Eg5 to cross-links microtubules thereby facilitating spindle assembly and mediating the sliding of antiparallel interpolar microtubules to increase metaphase spindle length. Also more importantly, NuSAP interacts with Kif2A on spindle microtubules in an Aurora A phosphorylation-dependent manner, thereby maintaining a proper amount of Kif2A at spindle poles through a phosphorylation and dephosphorylation cycle. Once the minus-ends of MTs are incorporated into spindle poles, Eg5 and Kif2A induce the proper MT poleward movement. When NuSAP is depleted in an experimental condition, the normal localization of Eg5 on spindle microtubules is reduced and excess Kif2A concentrates on spindle poles, leading to instability of MTs, and increased MT flux rate, and thus to a short metaphase spindle formation with chromosomal misalignment.

Our results showed Aurora A phosphorylates NuSAP at mitosis on a conserved serine residue S240. The affinity of NuSAP to Kif2A is S240 phospho-dependent, yet this is not the case for the NuSAP-Eg5 interaction. NuSAP is mainly localized on the spindle MTs near chromosomes. Thus, NuSAP and Kif2A may form complexes on the mitotic spindle microtubules. It is likely that through an unknown mechanism, conformational change possibly, phosphorylated NuSAP interacts with more Kif2A on the spindle body and then reduces the amount of Kif2A on spindle poles, thus restraining the excess accumulation of Kif2A at the minus ends of MTs to ensure the stability of mitotic spindles. Moreover, although NuSAP is responsible for recruiting Eg5 onto spindle MTs, the affinity of NuSAP to Eg5 is not S240 phospho-dependent. It has been reported that Eg5 activity was shown to be required for flux in *Xenopus* extract spindles and driving the spindle scale through exerting forces between antiparallel MTs (Miyamoto et al., 2004; Shirasu-Hiza, Perlman, Wittmann, Karsenti, & Mitchison, 2004). Biochemical depletion of Eg5 significantly decreases the flux rate(Miyamoto et al., 2004), which is opposite to our observations after NuSAP knockout. Thus, the increased MT rate induced by NuSAP knockout is mainly because of Kif2A but not Eg5. It remains to be determined how NuSAP balances the Kif2A-mediated microtubule depolymerization near the poles and the continuous antiparallel microtubule sliding caused by Eg5.

In conclusion, our findings shed light on the crucial roles of NuSAP in regulating mitotic spindle assembly, structural dynamics and metaphase spindle length control for proper chromosome alignment through modulating microtubule flux, in concert with Eg5, Kif2A and other regulators of the spindle MT dynamics.

## Materials and methods

**Key resource table.**
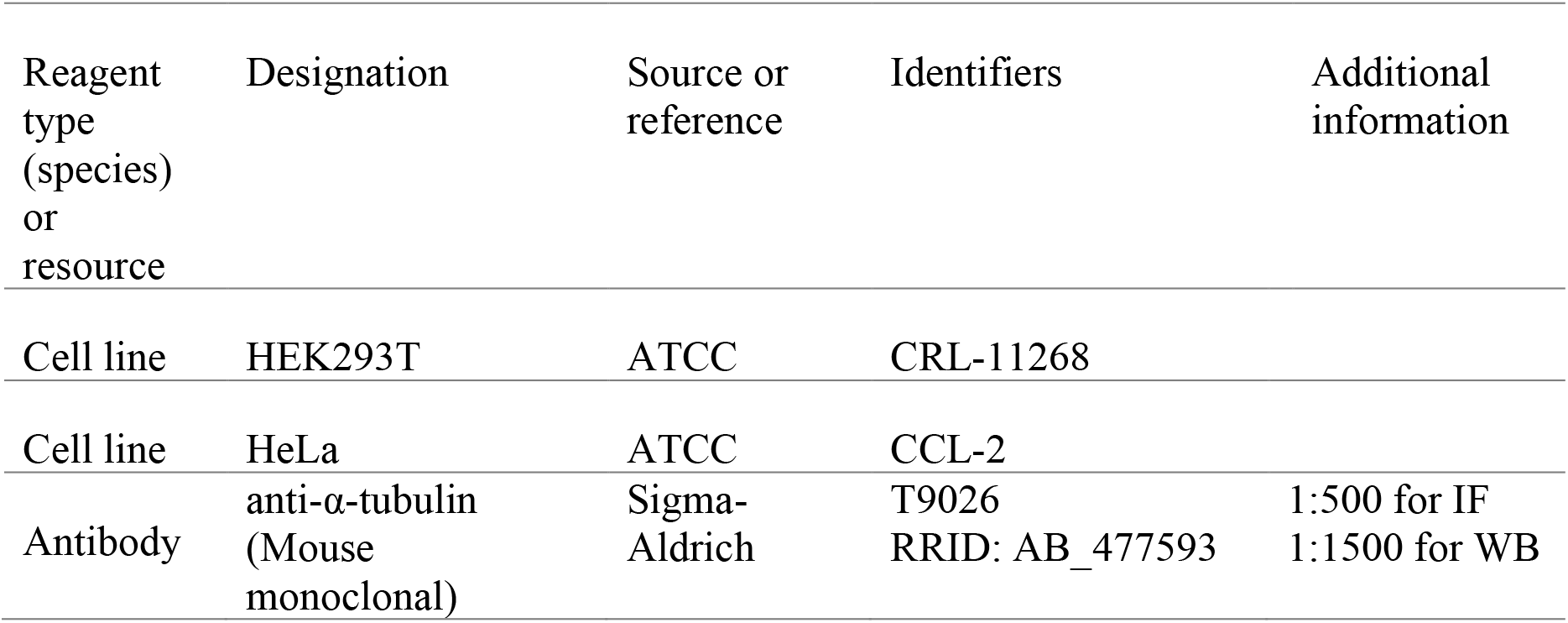
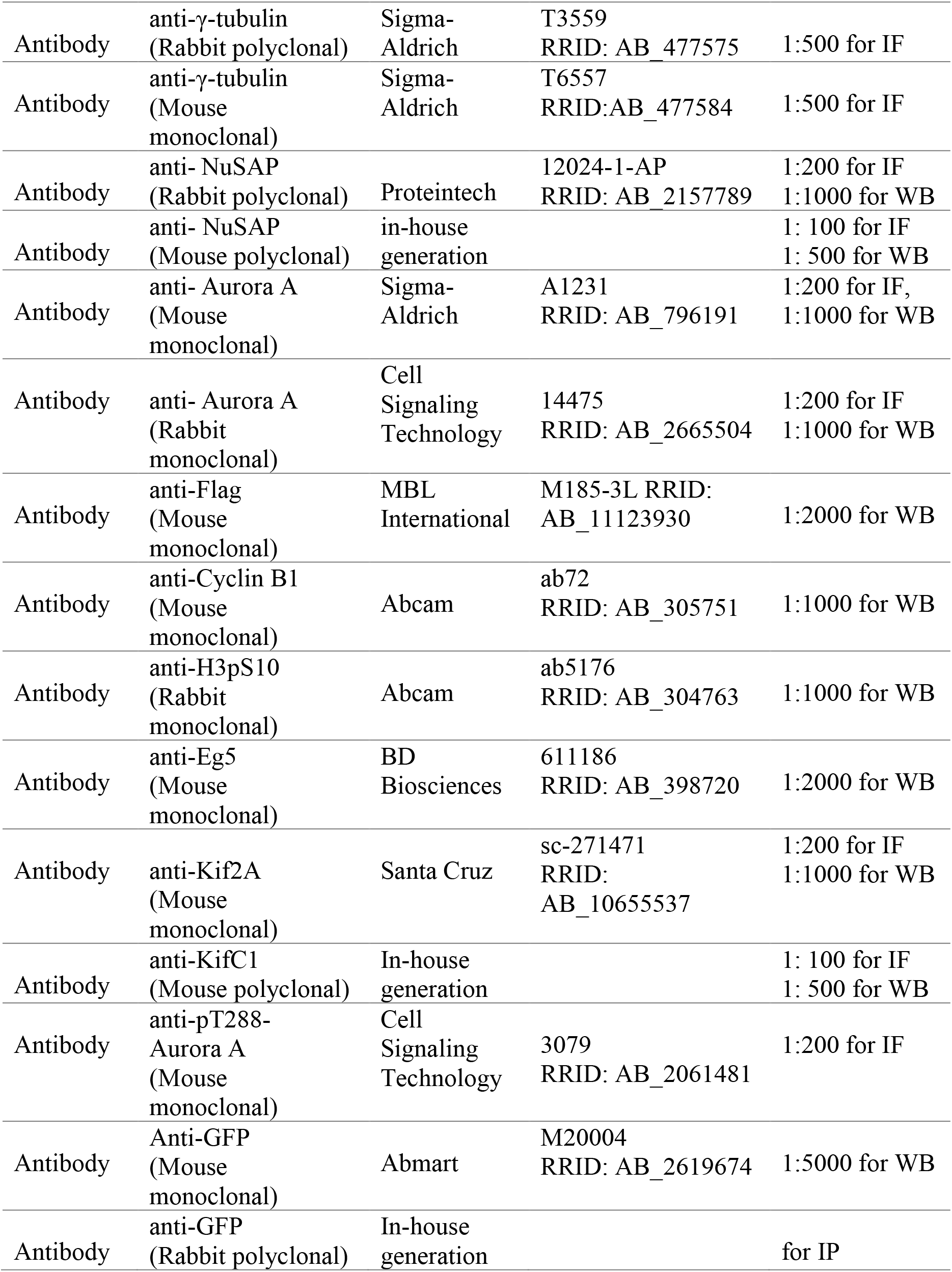
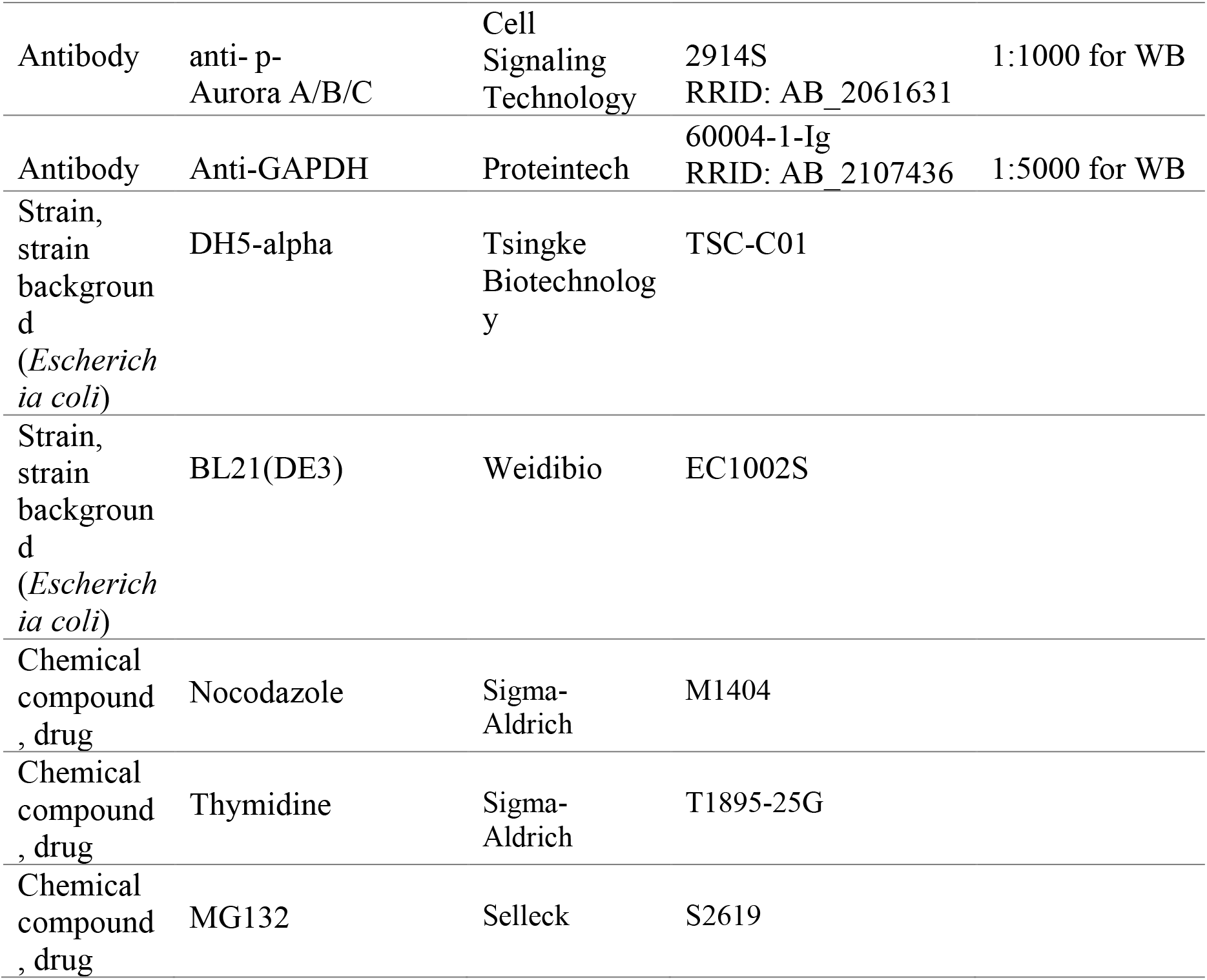

### Protein purification

GST-tagged proteins were affinity-purified using glutathione-Sepharose 4B beads (GE Healthcare) according to the manufacturer’s protocols. NuSAP polyclonal antibody was obtained by immunizing mice (Balb/c) with the full-length recombinant GST-tagged NuSAP. The anti-GFP antibody used for immunoprecipitation was generated by immunizing rabbits with bacterially expressed recombinant His-tagged GFP. All animal experiments were performed in the Laboratory Animal Center of Peking University in accordance with the National Institute of Health Guide for Care and Use of Laboratory Animals according to guidelines approved by the Institutional Animal Care and Use Committee at Peking University.

### Cell culture, synchronization, and transfection

HeLa and HEK293T cells were cultured in DMEM (CORNING) supplemented with 10% bovine calf serum (CellMax) at 37°C in a 5% CO2 incubator. For the double thymidine block and release experiment, the cells were arrested for 17-20 h with 2.5 mM thymidine (Sigma-Aldrich) with a 9 h release interval. After release into fresh medium, cells were harvested at the indicated time points. Mitosis-arrested cells were obtained by adding 100 ng/mL nocodazole (Sigma-Aldrich) for 12-15 h after release from the thymidine block. To synchronize to metaphase for immunofluorescence analysis, HeLa cells were treated with thymidine for 16-20 h, released for 9 h, and then incubated with 10 μM MG132 for 1 h to fully establish the bipolar spindle. Transient cDNA and siRNA transfections were performed with Lipofectamine^™^ 2000 (Invitrogen) according to the manufacturer’s instructions. The following siRNA sequence was used: NuSAP, 5’-GCACCAAGAAGCUGAGAAUTT-3’ (Raemaekers et al., 2003).

### CRISPR/Cas9-mediated editing of NuSAP genes in HeLa cells

The single-guide RNAs (sgRNAs) for targeting human NUSAP genes were predicted by CHOPCHOP (https://chopchop.cbu.uib.no/) (Ran et al., 2013). To express sgRNA (5’-CTGCAGGTCACTGTACTTGA-3’) for human NUSAP gene knockout, the oligonucleotides were annealed and cloned into the BbsI site of dual Cas9 and sgRNA expression vector pSpCas9(BB)-2A-Puromycin (Dr. Feng Zhang laboratory, Addgene, #48139). The plasmids were transfected into HeLa cells using TurboFect Transfection Reagent (Invitrogen). After 48h, cells were treated with 1 μg/ml puromycin for 2–3 days. Then the live cells were split individually to make a clonal cell line.

Clones with loss of NuSAP proteins were isolated by immunostaining and confirmed by immunoblotting. The genomic DNA PCR fragments were subcloned into pClone007 vector (Tsingke, pClone007 Blunt simple vector kit) and transformed into the *E. coli*. Then, certain numbers of bacterial colonies were sequenced to confirm the gene disruption. The PCR primers were as follows: forward 5’-ACTCGTTACCTGAACAGGCG-3’ and reverse 5’-TTCGTAATCGACGCCCTGAG-3’.

### Immunoprecipitation (IP) and Western blotting

Mitotic Cells were washed with cold PBS and lysed in cell lysis buffer (20 mM Tris-HCl, pH=8.0, 150 mM NaCl, 2 mM EGTA, 0.5 mM EDTA, 0.5% NP-40, 5 mM NaF, 1 mM Na3VO4, 1 mM PMSF, and 500× protease inhibitor cocktail; Calbiochem) for 20 min on ice. Lysates were centrifuged at 13,000 rpm for 15 min, and supernatants were incubated with For NuSAP IP assay, HeLa cell lysates were mixed with mouse anti-NuSAP polyclonal antibody (provided by Dr. Zhaoxuan Deng) and rabbit anti-NuSAP polyclonal antibody (Proteintech, 12024-1-AP). For anti-Flag and anti-GFP IP assays, HEK293T cells were transfected with the indicated constructs Flag-Aurora A and GFP or GFP-NuSAP, and the cell lysates were mixed with protein A–Sepharose beads (75% slurry) conjugated with indicated antibodies for 1 h at 4°C. After five washes with lysis buffer, the beads were suspended in Laemmli sample buffer. After being resolved on SDS-PAGE gels, the proteins were transferred to nitrocellulose membranes that were then blocked in TTBS (20 mM Tris-HCl, pH=7.5, 500 mM NaCl, and 0.3% Tween 20) containing 3% nonfat milk at room temperature for 1 h. Then, they were probed with primary antibodies (diluted in nonfat milk) and then with HRP-conjugated secondary antibody. The membranes were developed for visualization by enhanced chemiluminescence (Sigma-Aldrich) and X-ray film. The protein-protein interaction discrepancy was quantified by ImageJ. The relative intensity between mCherry-tagged NuSAP constructs and GFP-Kif2A or GFP-Eg5 was defined by the ratio of the amount of interacting mCherry-NuSAP proteins to that of immunoprecipitated GFP-Kif2A or GFP-Eg5. The value of the WT group was set as 1.0, and the other mutant groups were calculated.

### Immunofluorescence

Cells were grown on coverslips and fixed in precooled methanol for 5 min on ice followed by incubation with primary antibodies (diluted in 3% BSA) at 4°C overnight. After washing three times with PBS, the cells were incubated with fluorescence-labeled secondary antibodies for 1 h at room temperature. Coverslips were mounted with Mowiol (Sigma-Aldrich) containing 1 μg/mL DAPI (Sigma-Aldrich) for DNA staining. Images were acquired using an imaging system (DeltaVision; Applied Precision) equipped with an inverted microscope (IX-71; Olympus) and a 100×/1.42 NA oil objective. All immunofluorescence images were captured with a 4-μm Z-section thickness by 8-slices and processed for maximum intensity projection. The images were analyzed using Volocity (ver. 6.1.1) software.

### GFP-α-tubulin photoactivation analysis

For activation studies, mitotic HeLa cells transfected with photoactivatable GFP-α-tubulin and indicated mCherry-tagged constructs were identified by DIC microscopy and after the acquisition of three pre-activation frames, a 405 nm laser was used to activate GFP-α-tubulin in a rectangular region near the MT plus ends inside the spindle. Imaging was performed with a Spinning disc confocal microscope equipped with an inverted microscope (Nikon TiE) and a 60X/1.4 NA oil objective lens and images were acquired every 10 s. For MT poleward flux experiments, the quantification of fluorescence intensity of the activated areas and the distance of fluorescent signal movement were analyzed using Volocity software.

### λ protein phosphatase (λ-PPase) Assay

Mitotic HeLa cells were collected from a 35-mm dish and lysed with lysis buffer, containing 20 mM Tris-HCl, pH=8.0, 150 mM NaCl, 2 mM EGTA, 0.5% NP-40, and complete EDTA-free protease inhibitors (Roche) on ice for 20 min. Lysates were centrifuged at 13,000 rpm for 15 min, and supernatants were incubated with 10× NEBuffer for PMP, 10× 10 mM MnCl2, and 400 U λ-PPase (New England Biolabs, Inc.) at 30°C for 30 min before being stopped with SDS sample buffer. Then the reaction system was analyzed by Western blotting analysis.

### *In vitro* kinase assays

2 μg GST-tagged NuSAP point mutants were incubated with 50 ng human recombinant Aurora A (Sigma-Aldrich) in a water bath at 30°C for 30 min in kinase buffer (50 mM Tris-HCl, pH=7.5, 10 mM MgCl_2_, 2 mM EGTA, 5 mM DTT) containing 100 μM ATP, and 6,000 Ci/mmol γ-[^32^P] ATP (GE Healthcare). Reactions were quenched by the addition of Laemmli sample buffer and analyzed by SDS-PAGE and autoradiography.

### Measurement of relative fluorescence intensity

For fluorescence intensity of Kif2A on spindle poles, four square boxes of 3 x 3 μm were drawn, two on two spindle poles and the other two on their adjacent spindle areas. The mean fluorescence intensity was measured using Velocity software (version 6.1.1). The relative fluorescence intensity was defined by the ratio of the mean intensity on spindle poles and on the adjacent spindle areas. The relative fluorescence intensity of Eg5 (or KifC1) was defined by the ratio of the mean fluorescence intensity of Eg5 (or KifC1) and α-tubulin on the entire spindle microtubule area. The relative fluorescence intensity of MCAK was defined by the ratio of the mean fluorescence intensity of MCAK and Crest on the entire chromosome area.

### Statistical analysis

Statistical significance was performed using Graphpad Prism (ver. 6.0c) software. We did not pre-establish exclusion criteria. *P*-values were calculated with two-tailed *t* test from the mean values of the indicated data. All statistical data were presented as mean ± SD. Significant difference were marked with asterisks (**P<0.05; ***P<0.001; ****P<0.0001; ns, no significant difference).

## Acknowledgments

We thank Dr. Jiahuai Han (University of Xiamen) for providing *NUSAP1* cDNA and the National Center for Protein Sciences at Peking University (Beijing, China), particularly Drs. Hongxia Lv and Xiaochen Li, for technical help. We thank Dr. Zhaoxuan Deng for initiating this work and all other members of our laboratory for their constructive suggestions.

## Additional information

### Funding

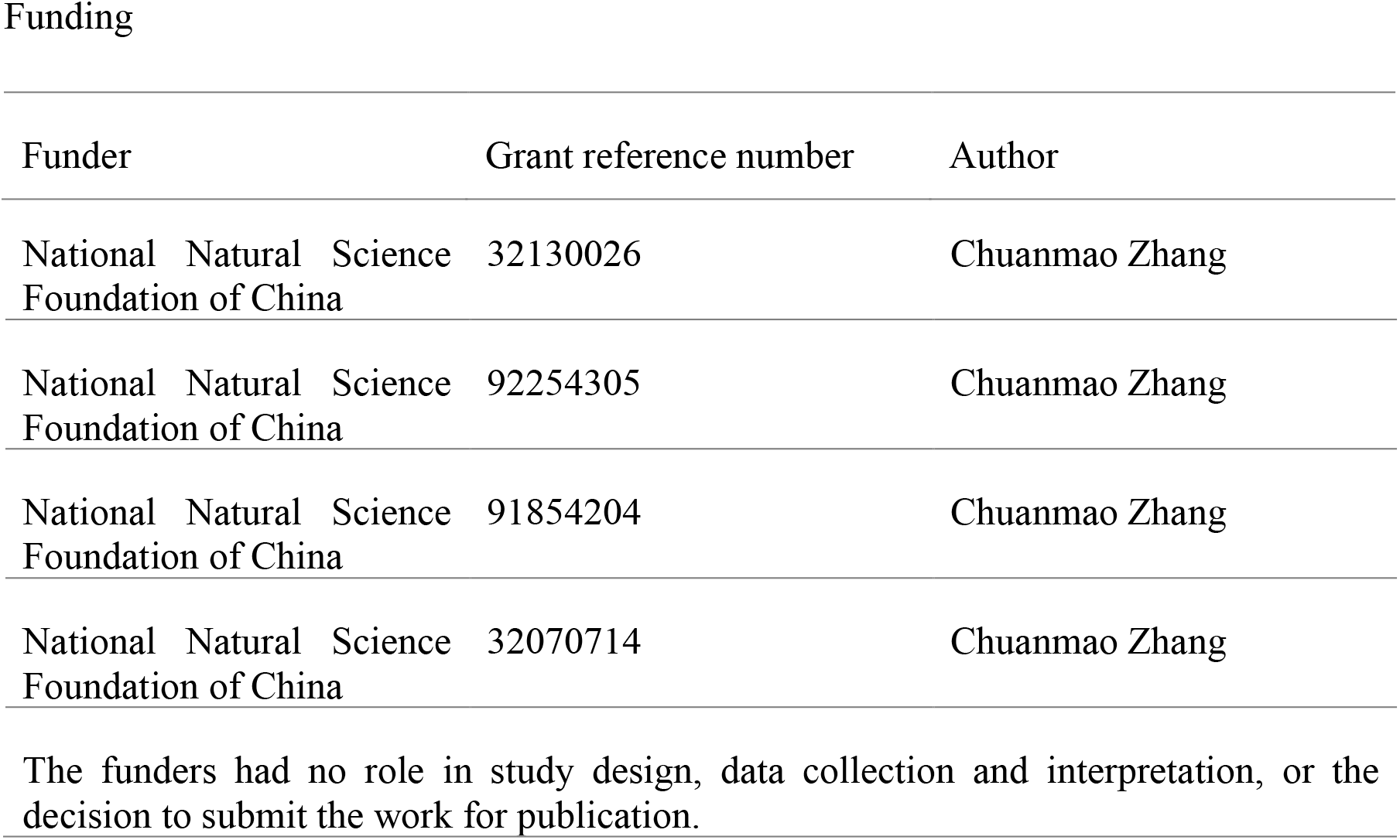

| Funder | Grant reference number | Author |
| --- | --- | --- |
| National Natural Science Foundation of China | 32130026 | Chuanmao Zhang |
| National Natural Science Foundation of China | 92254305 | Chuanmao Zhang |
| National Natural Science Foundation of China | 91854204 | Chuanmao Zhang |
| National Natural Science Foundation of China | 32070714 | Chuanmao Zhang |

The funders had no role in study design, data collection and interpretation, or the decision to submit the work for publication.

### Author contributions

CMZ conceived the project; CMZ, YW, MJS, GWX and QJ designed the experiments; YW, MJS, and GWX analyzed the data and performed most of the experiments; BYY, performed some of the experiments; and CMZ, YW, MJS, GWX and QJ wrote the manuscript. #These authors contribute equally to this paper.

### Data availability

All data generated or analyzed during this study are included in the manuscript and supporting files. Source data files have been provided for Figures 1–7 and figure supplements.

### Materials availability statement

A list of the reagents and newly created materials included in this study are available on request from the corresponding author.

---

**Figure 2-supplement 1.**
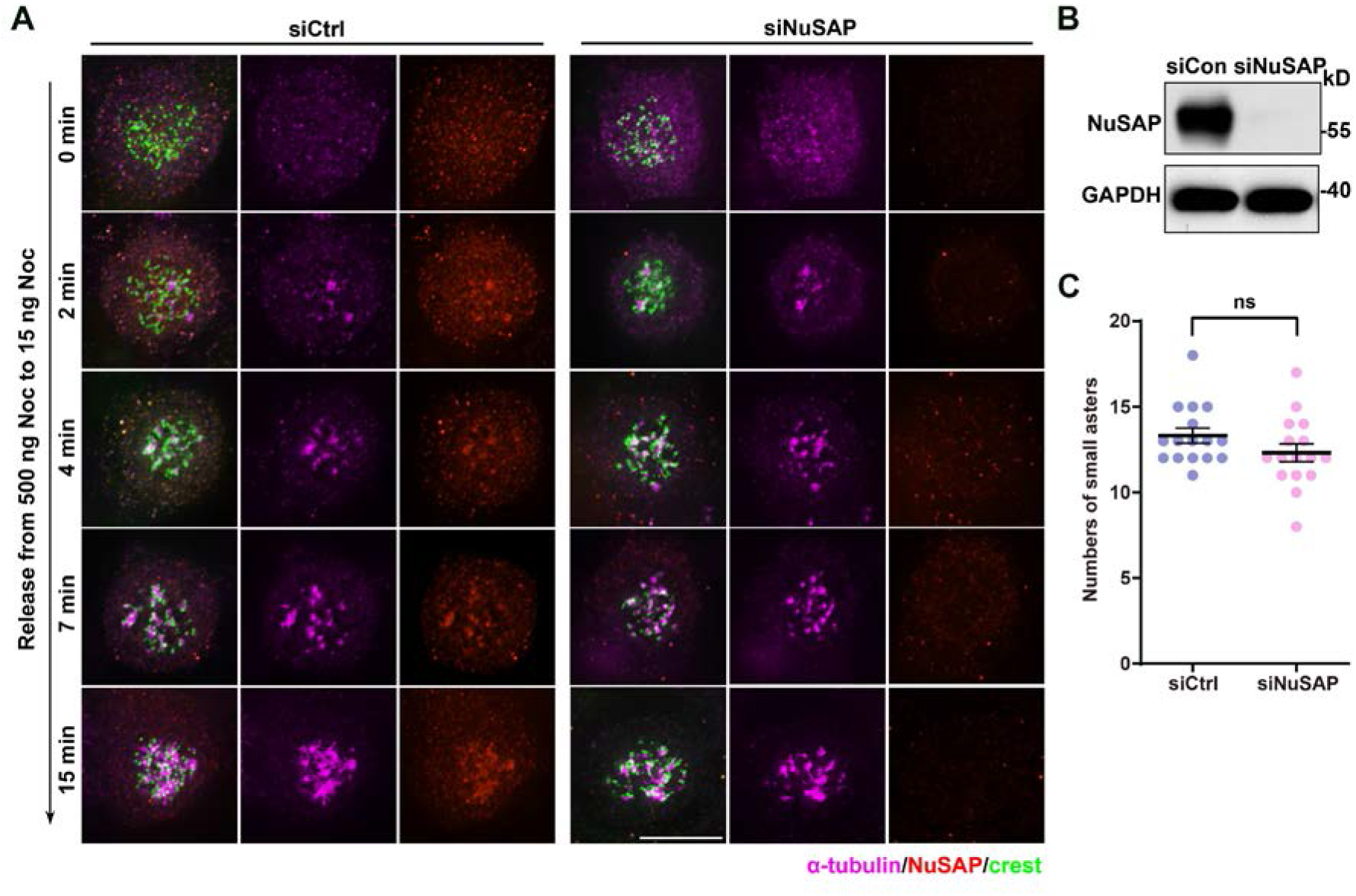
NuSAP is not necessary for acentrosomal MT nucleation. (A) HeLa cells were transfected with the indicated siRNAs and treated with 500 ng/ml nocodazole for 2 h. Then, the treated cells were released into medium containing 15 ng/ml nocodazole and fixed at the indicated time. (B) Detection of NuSAP RNAi knockdown efficiency in HeLa cells by Western blotting. (C) Quantification of the number of small acentrosomal MT asters when cells were released at the time point of 15 min (shown in A). Error bars indicated SD. Sixteen cells were quantified. Unpaired two-tailed t test. P=0.1516. ns, not significant. Scale bars, 10 μm. Figure 2-supplement 1-source data 1 Raw Microsoft excel file used for analysis of graphs in Figure 2-supplement 1C Figure 2-supplement 1-source data 2 Labeled uncropped western blot images and raw western blot images in Figure 2-supplement 1B.

**Figure 5-supplement 1.**
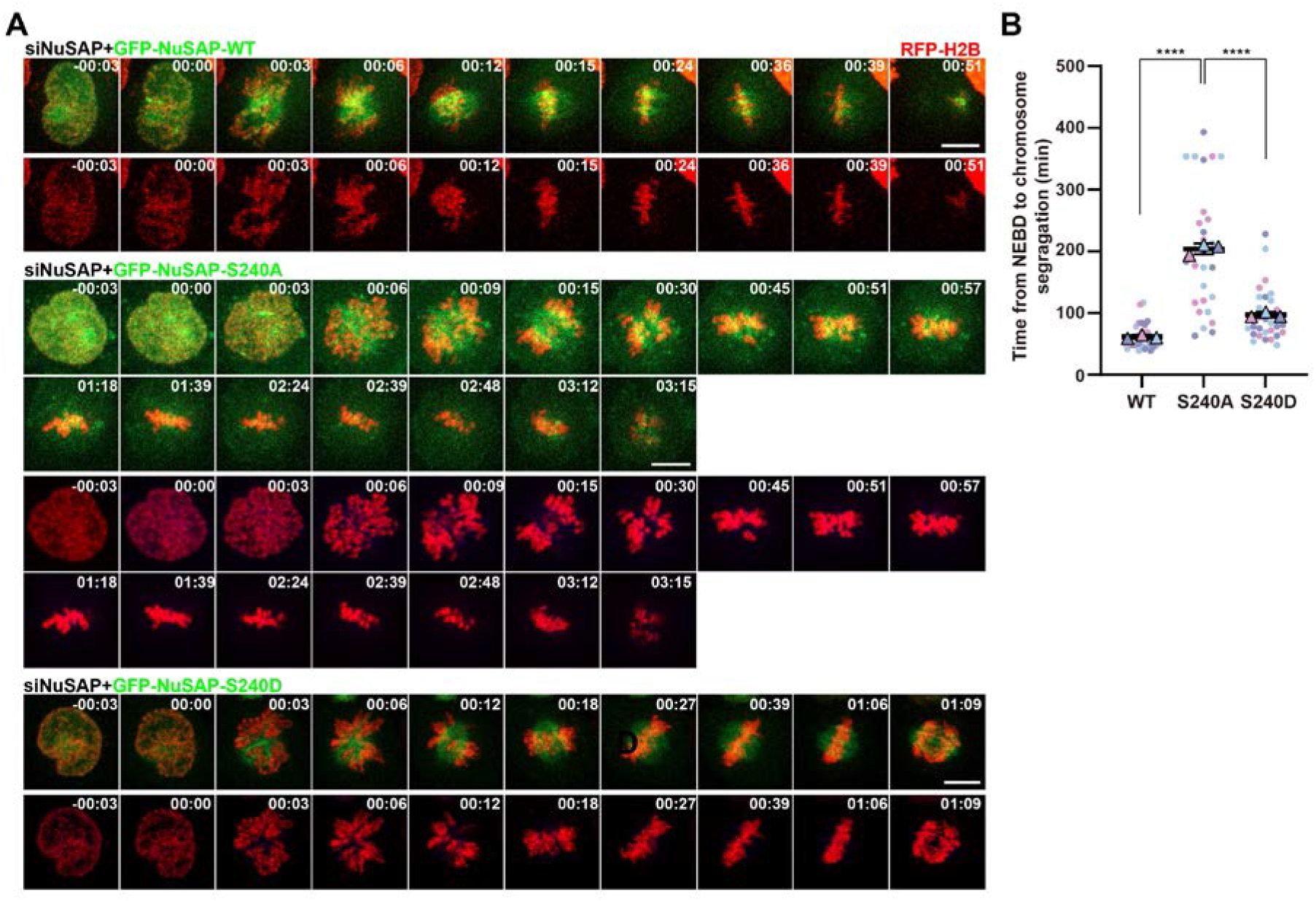
Phospho-null NuSAP resulted in abnormal spindle morphology and delayed mitosis progression. (A) RFP-H2B expressing HeLa cells with NuSAP siRNA knockdown were processed for rescue experiments with GFP-tagged NuSAP-WT, -S240A, and -S240D followed by live-cell imaging. Images were captured every 3 min with 5 slices of 8 μm Z-section thickness in total. (B) Quantification of the time from NEBD to chromosome separation shown in A. Error bars indicated SD. Three independent replicates of 10 cells per replicate were quantified. Unpaired two-tailed t test. P<0.0001 for WT/S240A. P<0.0001 for S240A/S240D. ****, P < 0.0001. Scale bars, 10 μm. Figure 5-supplement 1-source data 1 Raw Microsoft excel file used for analysis of graphs in Figure 5-supplement 1B.

**Figure 6-supplement 1.**
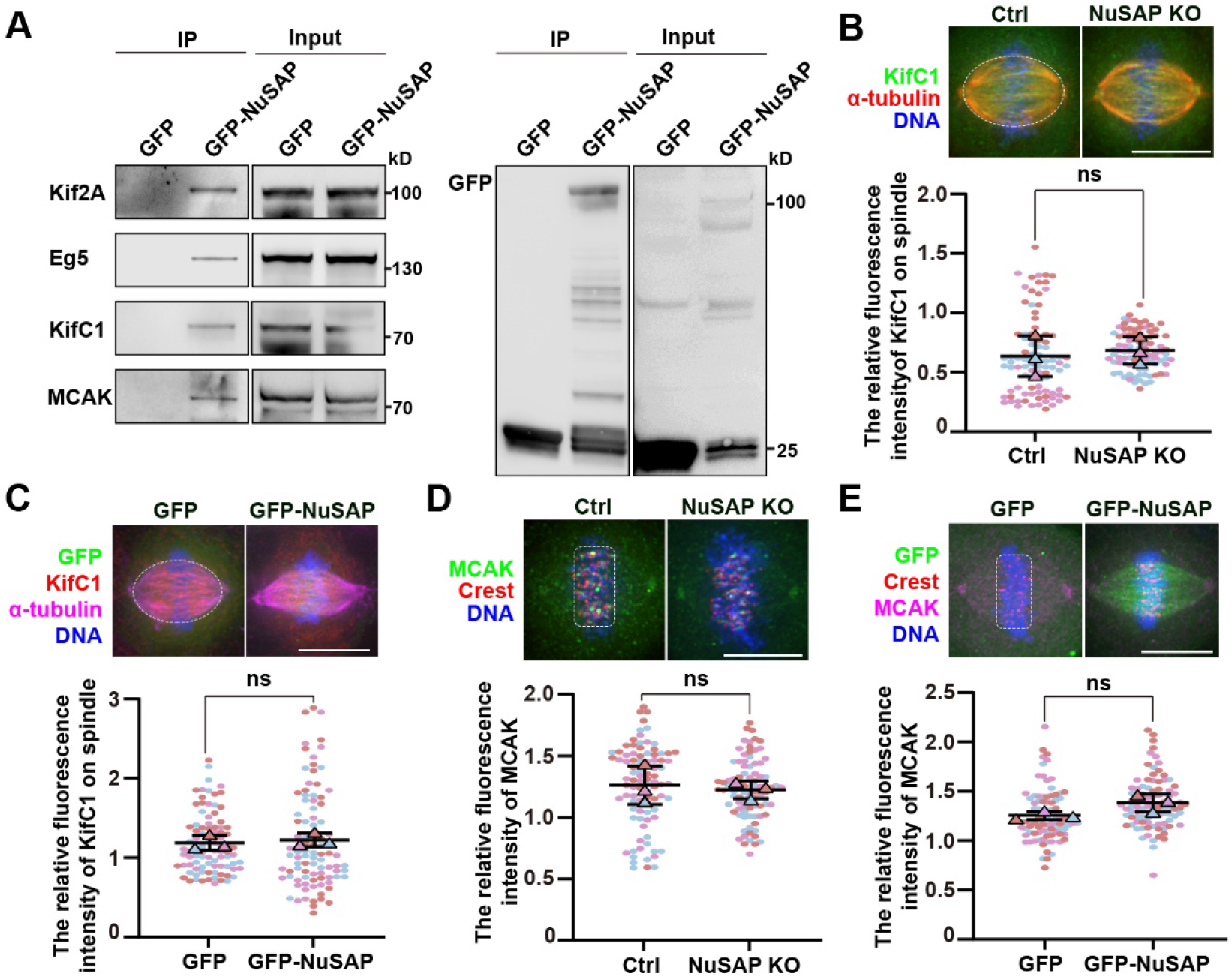
NuSAP had no effect on the localization of KifC1 and MCAK. (A) NuSAP interacted with Kif2A, Eg5, KifC1, and MCAK. Mitotic HEK293T cells were transfected with GFP or GFP-NuSAP and then immunoprecipitated with GFP-Trap beads. (B and D) Sample images (top) and quantification (bottom) of relative KifC1 (B) and MCAK (D) fluorescence intensity on spindle of control and NuSAP-knockout cells, respectively. Error bars indicated SD. Three independent replicates of 30 cells per replicate were quantified. Unpaired two-tailed t test: P = 0.7074 in (B); P = 0.7179 in (D). The white dotted lines show the areas of measurement.(C and E) Sample images and quantification of relative KifC1 (C) and MCAK (E) fluorescence intensity on spindle of HeLa cells transfected with GFP or GFP-NuSAP, respectively. Error bars indicated SD. Three independent replicates of 30 cells per replicate were quantified. Unpaired two-tailed t test: P = 0.6413 in (C); P = 0.0880 in (E). The white dotted lines show the areas of measurement. Figure 6-supplement 1-source data 1 Raw Microsoft excel file used for analysis of graphs in Figure 6-supplement 1B-E. Figure 6-supplement 1-source data 2 Labeled uncropped western blot images and raw western blot images in Figure 6-supplement 1A.

## Videos with still images and legends

**Video 1.**
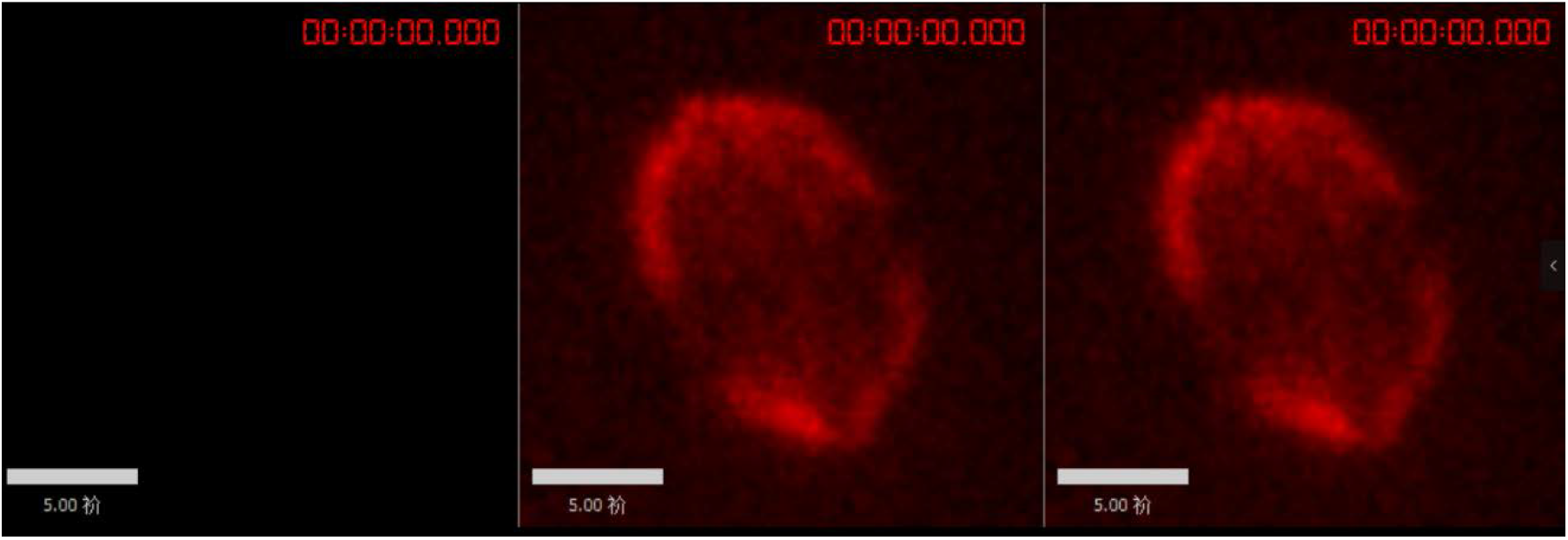
MT flux on metaphase spindles in HeLa cells. Control HeLa cells were transfected with photoactivatable GFP-tagged α-tubulin (PAGFP-α-tubulin). 405 nm laser was applied to activate GFP signal in a rectangular region near the MT plus ends, and the movement of the fluorescent mark (green) was tracked every 10 s. Microtubules were stained with SiR-tubulin Scale bar, 5 μm.

**Video 2.**
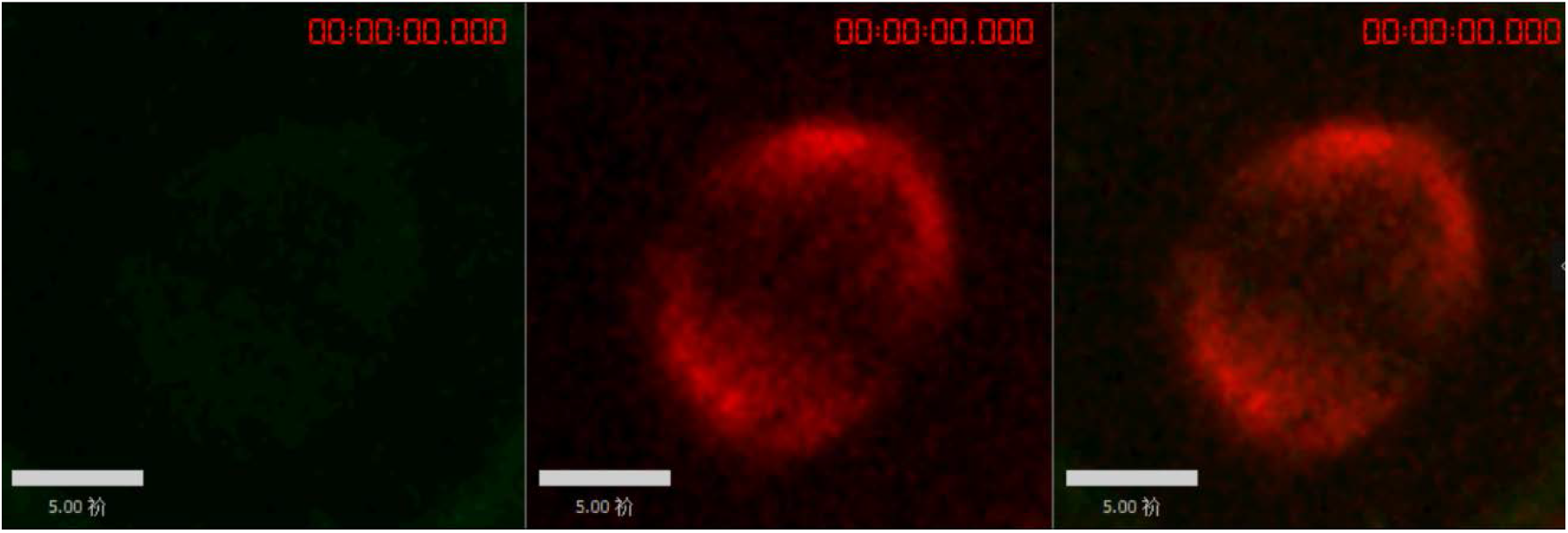
MT flux on metaphase spindles in HeLa cells with NuSAP knockout. NuSAP-knockout HeLa cells were transfected with photoactivatable GFP-tagged α-tubulin (PAGFP-α-tubulin). 405 nm laser was applied to activate GFP signal in a rectangular region near the MT plus ends, and the movement of the fluorescent mark (green) was tracked every 10 s. Scale bar, 5 μm.

**Video 3.**
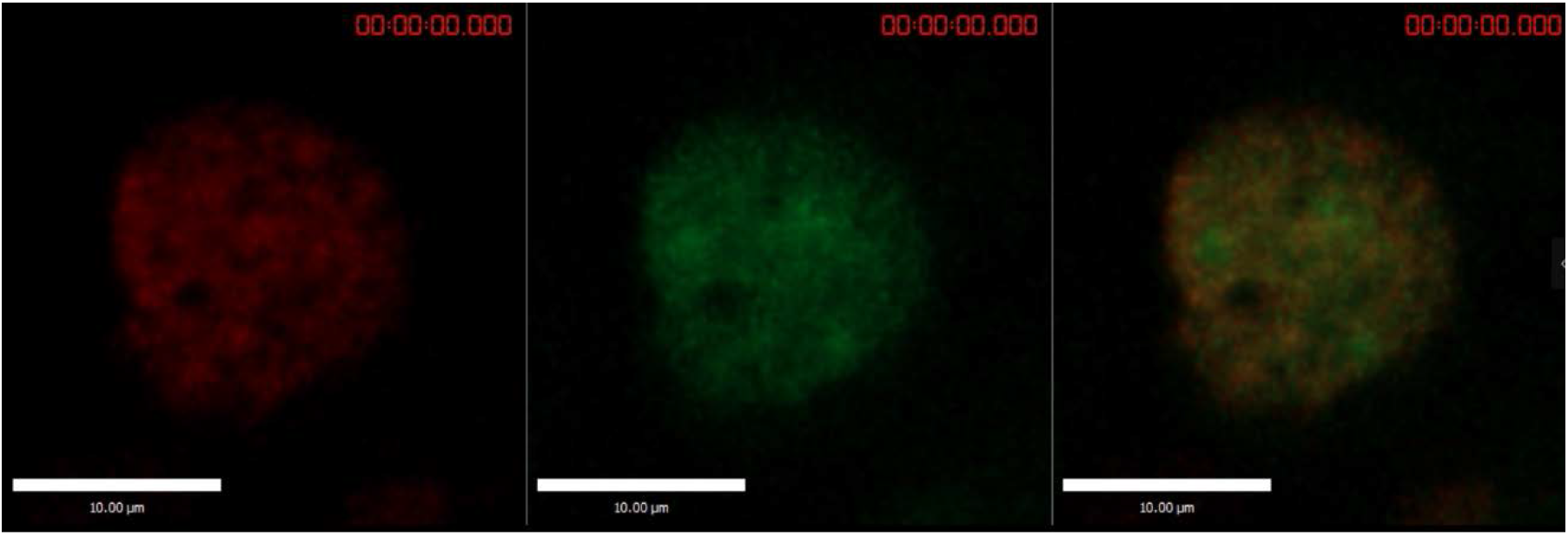
Spindle formation and chromosome alignment in the background of GFP-NuSAP-WT. HeLa cells stably expressing RFP-H2B (red) were depleted of endogenous NuSAP and transiently expressing RNAi-resistant GFP-NuSAP-WT (green). Images were captured every 3 minutes. Bar, 10 μm.

**Video 4.**
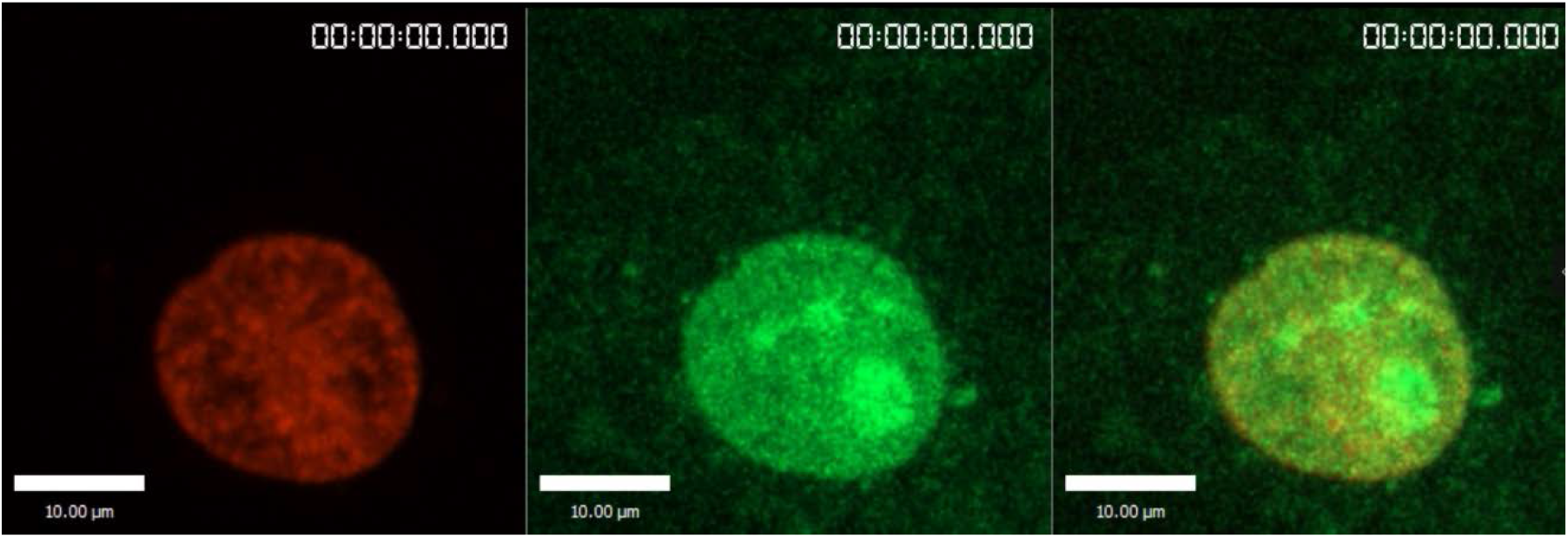
Spindle formation and chromosome alignment in the background of GFP-NuSAP-S240A. HeLa cells stably expressing RFP-H2B (red) were depleted of endogenous NuSAP and transiently expressing RNAi-resistant GFP-NuSAP-S240A (green). Images were captured every 3 minutes. Bar, 10 μm.

**Video 5.**
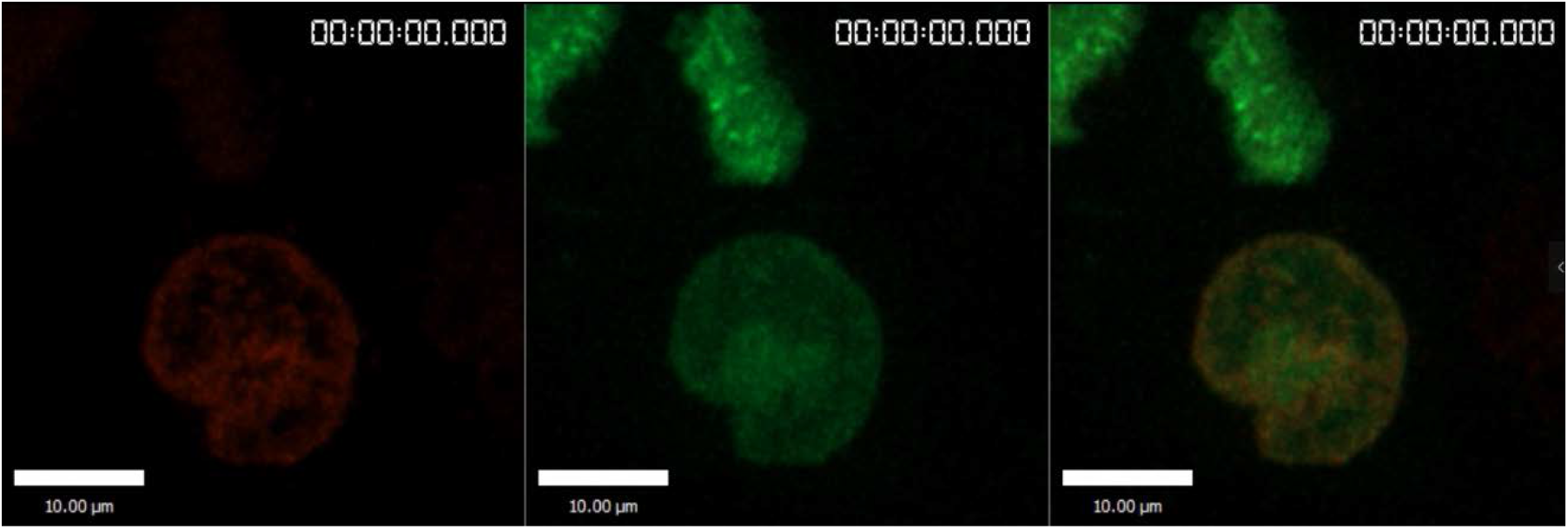
Spindle formation and chromosome alignment in the background of GFP-NuSAP-S240D. HeLa cells stably expressing RFP-H2B (red) were depleted of endogenous NuSAP and transiently expressing RNAi-resistant GFP-NuSAP-S240D (green). Images were captured every 3 minutes. Bar, 10 μm.

**Video 6.**
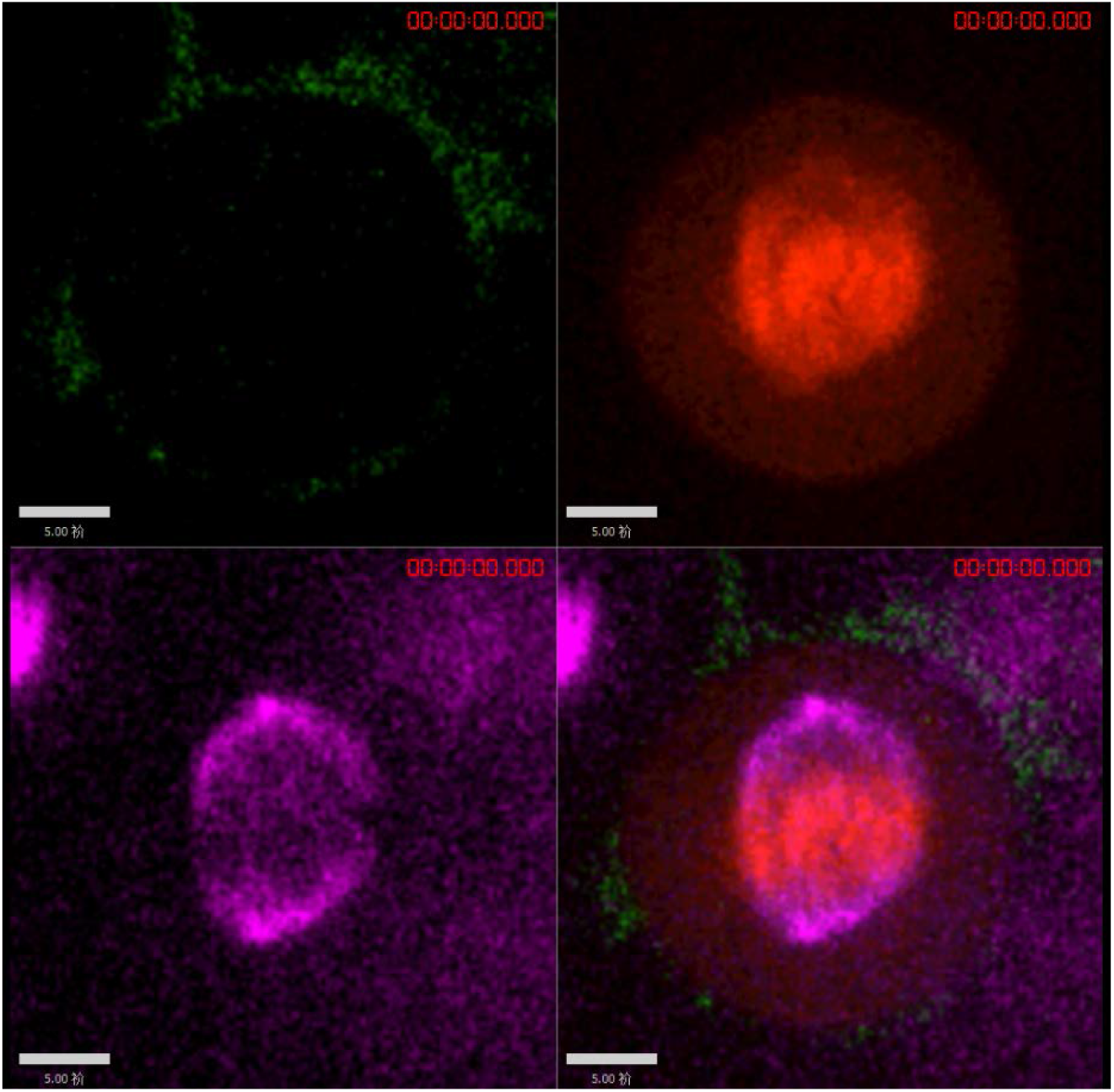
MT flux on metaphase spindles in HeLa cells in the background of mCherry-NuSAP-WT. NuSAP-knockout HeLa cells were co-transfected with photoactivatable GFP-tagged α-tubulin (PAGFP-α-tubulin) and mCherry-NuSAP-WT (red). 405 nm laser was applied to activate GFP signal in a rectangular region near the MT plus ends, and the movement of the fluorescent mark (green) was tracked every 10 s. Scale bar, 5 μm.

**Video 7.**
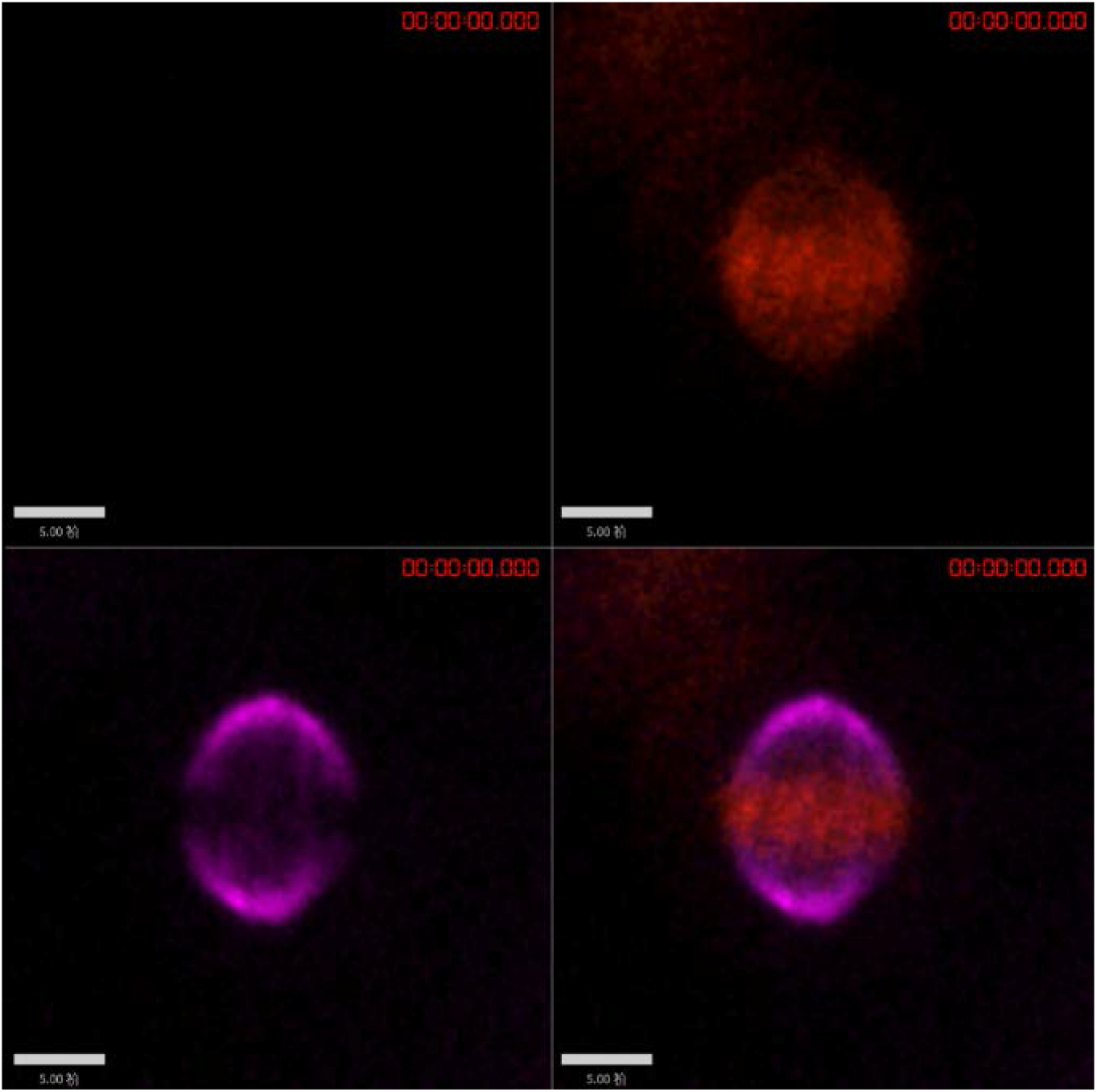
MT flux on metaphase spindles in HeLa cells in the background of mCherry-NuSAP-S240A. NuSAP-knockout HeLa cells were co-transfected with photoactivatable GFP-tagged α-tubulin (PAGFP-α-tubulin) and mCherry-NuSAP-S240A (red). 405 nm laser was applied to activate GFP signal in a rectangular region near the MT plus ends, and the movement of the fluorescent mark (green) was tracked every 10 s. Scale bar, 5 μm.

**Video 8.**
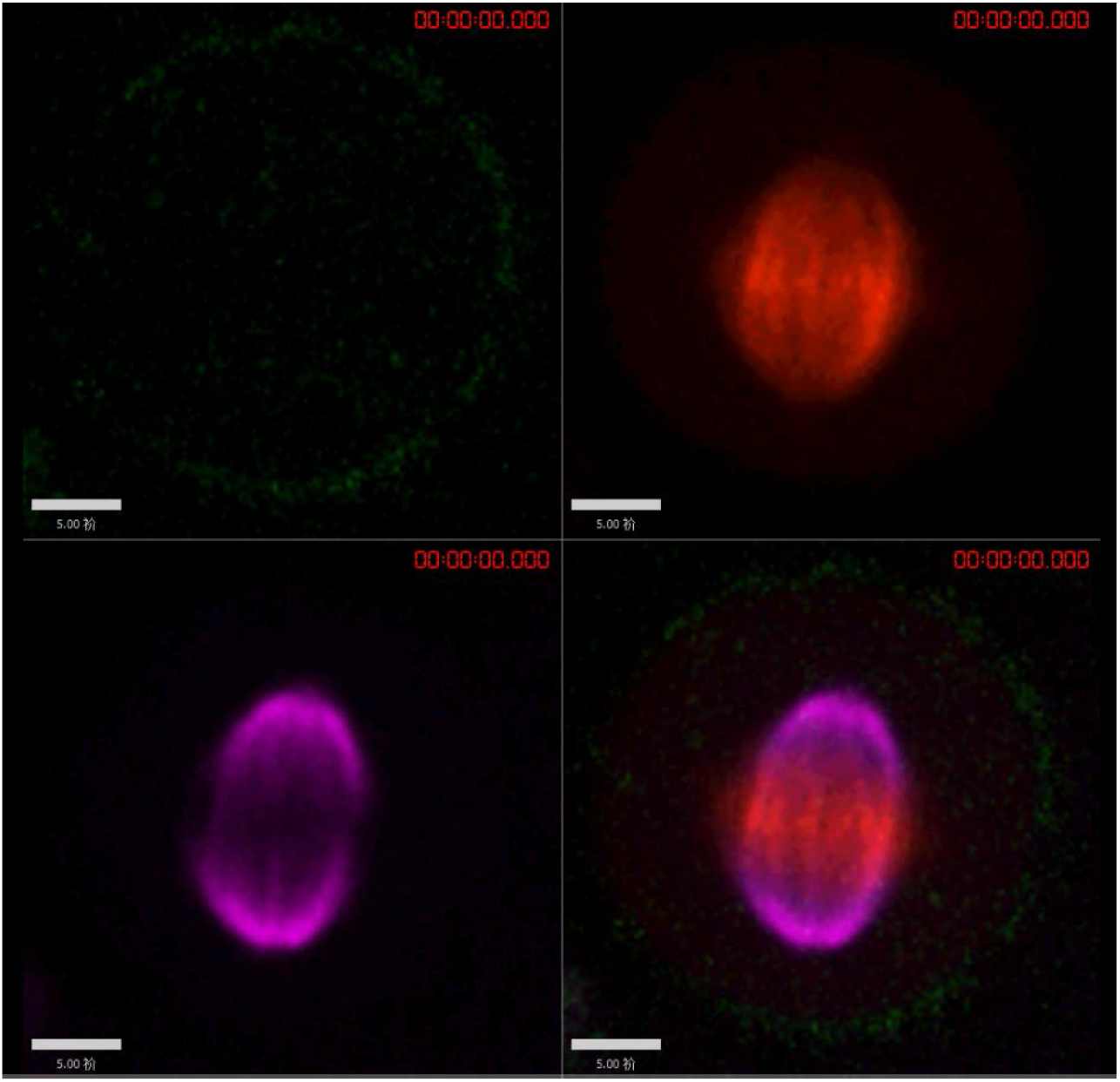
MT flux on metaphase spindles in HeLa cells in the background of mCherry-NuSAP-S240D. NuSAP-knockout HeLa cells were co-transfected with photoactivatable GFP-tagged α-tubulin (PAGFP-α-tubulin) and mCherry-NuSAP-S240D (red). 405 nm laser was applied to activate GFP signal in a rectangular region near the MT plus ends, and the movement of the fluorescent mark (green) was tracked every 10 s. Scale bar, 5 μm.

